# HTL/KAI2 signalling substitutes for light to control plant germination

**DOI:** 10.1101/2022.03.30.486460

**Authors:** Michael Bunsick, Zhenhua Xu, Gianni Pescetto, George Ly, Jenna Hountalas, François-Didier Boyer, Christopher S. P. McErlean, Julie D. Scholes, Shelley Lumba

## Abstract

Deciphering signalling pathways is essential to understanding how organisms respond to environmental cues but elucidating how these signalling pathways evolve in new environments is less clear.^1,2^ Most plants, for example, monitor multiple environmental cues to optimize the time and place to germinate. Some root parasitic plants, however, germinate in response to small molecules like strigolactones (SLs) emanating from host roots^3,4^ whilst a number of ephemeral weeds germinate in response to chemicals called karrikins (KARs) released after a forest fire.^5,6^ Although these species represent distinct clades, they use the same HYPOSENSITIVE TO LIGHT/KARRIKIN INSENSITIVE 2 (HTL/KAI2) signalling pathway to perceive strigolactones or karrikins, which suggests convergent evolution.^3,5^ Because specialist lifestyles are derived traits, it is not clear if HTL/KAI2 signalling in these species evolved from a specific germination-signalling pathway or whether this pathway had other functions that were co-opted for specialist germination circumstances. Here, we show HTL/KAI2 signalling in *Arabidopsis* bypasses the light requirement for germination. In part, this is because the HTL/KAI2 downstream component, SMAX1 impinges on PHYTOCHROME INTERACTING FACTOR 1/PHYTOCHROME INTERACTING FACTOR 3-LIKE 5 (PIF1/PIL5)-regulated hormone response pathways conducive to germination. We identified *Arabidopsis* accessions that can germinate in the dark, which had altered expression of HTL/KAI2 signalling components, suggesting that divergence in this signalling pathway occurs in nature. Moreover, *Arabidopsis HTL/KAI2*-regulated gene signatures were observed in germinating *Striga* seed. The ability of HTL/KAI2 signalling to substitute for light advances an explanation for how some specialist plants evolved their underground germination behaviour in response to specific environments.

## RESULT AND DISCUSSION

Parasitic plants like *Striga hermonthica* (*Striga*) and some pyroendemic plants like *Brassica tournefortii* germinate underground and do not require a light stimulus.^5,7,8^ In the lab, *Striga* and *B. tournefortii* seeds are typically germinated in the dark by adding SLs or KARs, respectively.^7,8^ Although SLs and KARs are distinct small molecules, they appear to signal through a class of related *α/β* hydrolase receptors called HTL/KAI2 in these species.^5,9,10^ In *Arabidopsis thaliana* (*Arabidopsis*), mutations in the HTL/KAI2 (AtKAI2) receptor were originally identified through genetic screens as positive regulators of hypocotyl light responses and KARs enhance the effect of light on germination and hypocotyl growth.^11,12,13^ These results suggested that limiting light conditions might uncover new roles for this receptor with respect to germination. Generally, *Arabidopsis* germination assays involve sterilizing seeds in the light and plating them onto nutrient rich agar. To create conditions more reflective of underground germination, we plated dry, unsterilized, and after-ripened seeds of the common laboratory wild type, Col-0, onto water agar plates and placed them directly in the dark. Imbibed Col-0 seeds did not germinate under these conditions but did show a slow and steady increase in germination over time under low light exposure (0.3 µE) (**Figure 1A**). A loss-of function At*KAI2* allele (*htl-3*) also did not germinate in the dark but low light exposure resulted in minimal germination suggesting At*KAI2* signalling contributes to light-dependent germination (**Figure 1A**). Three additional lines of evidence supported this premise. First, a previously characterized *Arabidopsis* line expressing the *Striga* ShHTL7 receptor (*ShHTL7OX*), which strongly activates the AtKAI2 signalling pathway,^14^ germinated in the dark on nanomolar levels of a racemic mixture of the artificial SL, *rac*-GR24 (**Figures 1B and 1C**). Second, genetic inactivation of two downstream regulators in KAI2 signalling,^15^ *SUPPRESSOR OF MAX2 1* (*SMAX1*) and *SUPPRESSOR OF MAX2 LIKE 2* (*SMXL2*), resulted in constitutive dark germination (**Figure 1B**). Finally, *htl-3* Col-0 lines misexpressing *KAI2* (*AtKAI2OX*) germinated in the absence of light at effective KAR_2_ concentrations (EC_50_) in the low nanomolar range (**Figures 1B and 1D**). Consistent with this, expression of genes (*DLK2, KUF1, BBOX20*) known to be induced by activation of AtKAI2^16^ increased in KAR_2_-treated AtKAI2OX seed and these genes were constitutively expressed in *smax1-2* single and *smax1-2 smxl2-1* double mutant (**Figure 1E**).

**Figure 1.**
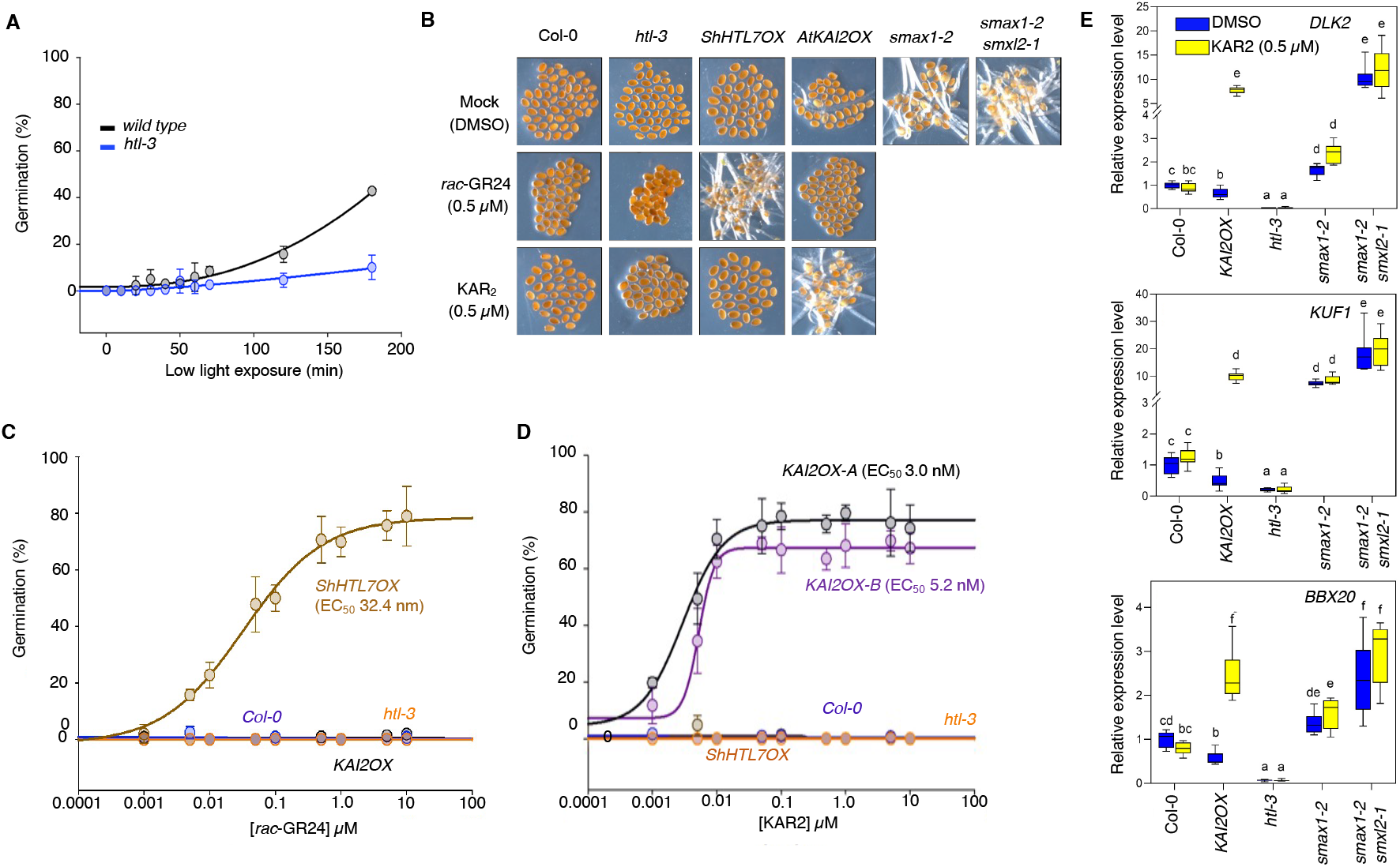
Dark germination in *Arabidopsis*. **(A)** Germination of Col-0 and an *AtKAI2* loss-of-function mutant (*htl-3*) seed in the dark and after discrete low light exposures (0.3 µE). Each point represents three independent biological replicates. Bar=SD (**B**) Dark germination of *Arabidopsis* genotypes placed on minimal media (mock or DMSO), 0.5 µM strigolactone (*rac*-GR24) or 0.5 µM karrikin (KAR_2_) containing media under dark conditions. (**C**) Effective concentrations (EC_50_) of *rac*-GR24 required to germinate 50% of Col-0, *htl-3* and an *ShHTL7OX* line. Each point represents three independent biological replicates. Bar=SD. (**D**) Effective concentrations (EC_50_) of KAR_2_ required to germinate 50% of Col-0, *htl-3, ShHTL7OX* and two *KAI2* misexpression lines *(KAI2OX-A, KAI2OX-B)*. Each point represents three independent biological replicates. Bar=SD. (**E**) Expression (RT–qPCR analysis) of KAR-induced genes (*DLK2, KUF1, BBOX20*) in dark-germinated *Arabidopsis* seeds on 0.5 µM KAR_2_. An 18S rRNA gene was used as an internal control. Letters indicate significant differences (*P* < 0.05, ANOVA with post-hoc Tukey HSD Test). *P* values, sample sizes and box plot elements are provided in **Table S1**.

The germination of *AtKAI2OX* seed at low nanomolar KAR_2_ concentrations in the dark is somewhat unexpected because adding KAR_2_ to *AtKAI2OX* seed in the light only partially rescued seed germination in seeds depleted for the germination stimulant, gibberellin (GA).^14^ It is thought this is because KAR_2_ does not fully activate the AtKAI2 signalling pathway. Possibly KAR_2_ is metabolized to a more potent ligand in the dark or light increases its degradation. It is also possible that dark is a less complex environment that sensitizes *Arabidopsis* germination to AtKAI2 signalling. Surprisingly, *AtKAI2OX* seed did not germinate in response to *rac*-GR24 in the dark although the 2’*S* enantiomer of this racemic mixture preferentially activates AtKAI2 in other assays^17^ (**Figure S1A**). When each enantiomer was tested separately, however, the 2’*S*-GR24 germinated At*KAI2OX* seed in the dark whilst 2’*R* isomer had no effect (**Figure S1B**). This suggested that the 2’*R* isomer was not only inactive but also inhibited 2’*S*-GR24 action in dark germination assays. Because *rac*-GR24 is active in many SL assays in the light, the clarity of enantiomeric specificity in dark germination may again reflect the simplicity of these conditions with respect to germination cues. Perhaps light establishes metabolic conditions that modify 2’*R* isomers thereby removing its inhibitory effects.

The transition from dark- to light-driven development (photomorphogenesis) requires light activated photoreceptors to inhibit specific negative regulators called *PHYTOCHROME INTERACTING FACTORS* (*PIFs)*.^18^ *PIF1* (also known as *PIL5* and BHLH105) is the major repressor of photomorphogenic germination and loss-of-function *pif1-1* mutant seed germinate in the dark.^19^ This *pif1* constitutive dark germination phenotype was epistatic to the decreased light sensitivity of *htl-3*, indicating that *PIF1* acts genetically at or downstream of *AtKAI2* (**Figure 2A**). Because of its repressive role in light responses, a simple model is PIF1 function requires SMAX1/SMXL2 action. Consistent with this, transcripts of direct targets of PIF1 repression, *HECATE 2* (*HEC2*) and *LONG HYPOCOTYL IN FAR RED 1* (*HFR1*),^20,21^ were highly induced upon activation of a *KAI2OX* line as were expression levels of photosynthetic genes (LHC, RBC, PS1 and PSII) (**Figure 2A**).

**Figure 2.**
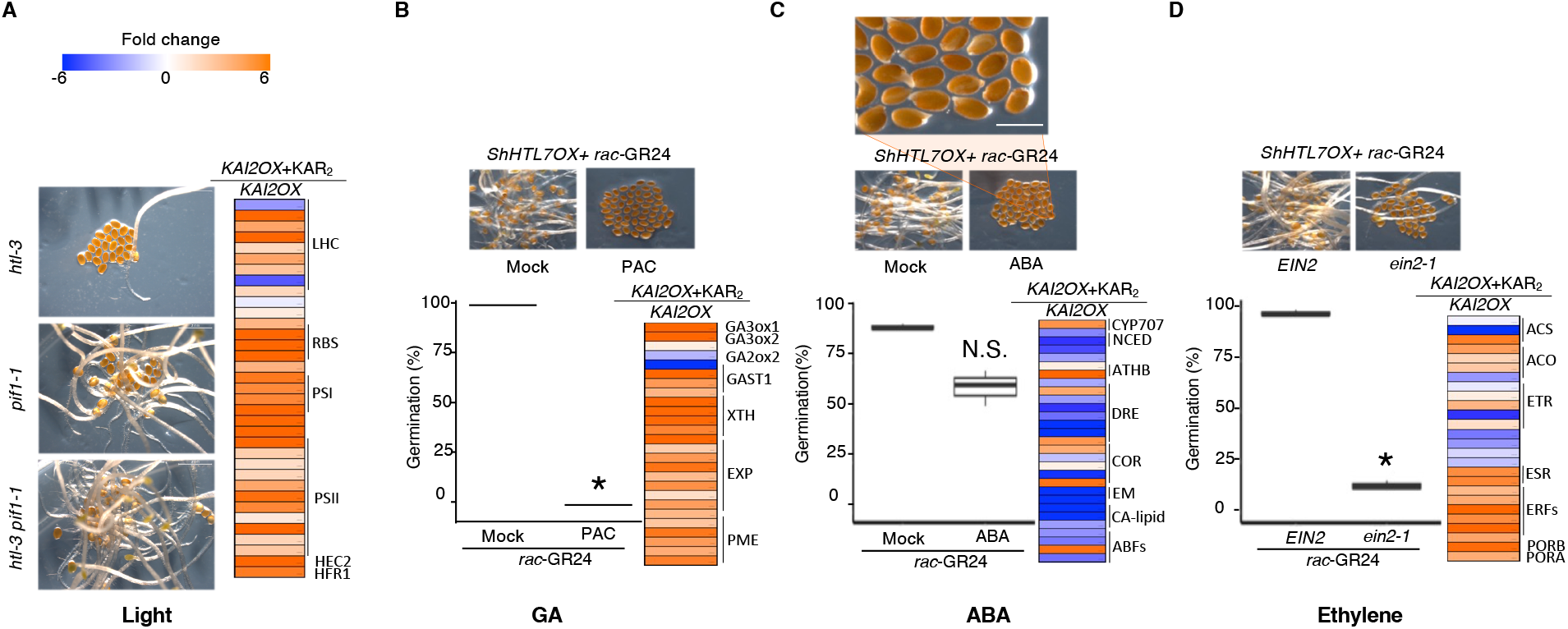
Dark germination of activating *AtKAI2* signalling lines perturbed in light or hormone signalling. (**A**) Left panel, representative pictures of *pif1-1, htl-3* and *pif1-1 htl-3* seed germinating in the dark. Right panel, heatmap of light-regulated genes expression in response to *rac*-GR24 addition in an *ShHTL7OX* line. (**B**) Germination of an *ShHTL7OX* line on paclobutrazol (PAC). Boxplot analysis represents three sets of biological replicates. Asterisk indicates a significant difference from the mock treatment (*P* < 0.05, ANOVA with one-sided Fisher’s LSD Test). Heatmap represents select GA-regulated genes in response to *rac*-GR24. (**C**) Germination of *ShHTL7OX* seed on ABA. Boxplot analysis represents three sets of biological replicates. Heatmap represents select ABA-regulated genes in response to *rac*-GR24. (**D**) Germination of *ShHTL7OX* seed containing a wild type *EIN2* or the loss-of-function *ein2-1* allele. Asterisk indicates a significant difference from *EIN2* (*P* < 0.05, ANOVA with one-sided Fisher LSD Test). Boxplot analysis represents three sets of biological replicates. Heatmap represents select ethylene-regulated genes in response to *rac*-GR24. The scale bar represents expression values on *rac*-GR24 relative to the control that have been log_2_-transformed. The heatmaps depicts fold-change at 24h. See **Table S2** for gene lists and values.

PIF1 also represses transcription of a seed specific regulator, SOMNUS (SOM), which influences gene expression of abscisic acid (ABA) and GA synthesis^22^. To functionally test the roles of these hormones on *AtKAI2* signalling, we fully activated the pathway by exposing *ShHTL7OX* seed to *rac*-GR24 and the addition of either ABA or paclobutazol (PAC), a GA biosynthetic inhibitor to decrease GA levels. Both conditions decreased the ability of *ShHTL7OX* seed to germinate (**Figure 2B, C**). Whilst PAC completely suppressed germination of *ShHTL7OX* seed on *rac*-GR24, the addition of ABA did not inhibit seed coat rupture but did inhibit further development (**Figure 2B, C**). This is consistent with observations that ABA alone cannot increase a DELLA repressor of GA signalling, REPRESSOR OF GA LIKE 2 (RGL2), to levels required to inhibit seed coat rupture.^23^ Consistent with the positive role of GA and negative role of ABA in germination, we found GA- and ABA-responsive genes increased and decreased respectively in KAR_2_-treated *AtKAI2OX* seed (**Figures 2B, C**). Moreover, we found *AtKAI2* signalling affected expression of SOM-regulated target genes involved in GA production (*GA3ox1, GA3ox2, GA2ox2*) and SOM-regulated genes involved in reducing ABA levels (*ABA1, NCED6, NCED9*) (**Figures 2B, 2C**).

Finally, when we introduced a mutation that confers ethylene insensitivity, *ein2-1* (*ETHYLENE INSENSITIVE 2*) into an *ShHTL7OX* line, it suppressed the dark germination phenotype (**Figure 2D**). Moreover, activation of *AtKAI2* increased transcript levels of genes involved in ethylene synthesis (*ACS7, ACO1, ACO2*) and affected expression of genes involved in ethylene action (**Figure 2D**). Interestingly, ethylene partially facilitates dark to light seedling transition by working in concert with *PIF1* to induce genes involved in chlorophyll synthesis.^24^ Consistently, both *PROTOCHLOROPHYLLIDE OXIDOREDUCTASE A* (*PORA*) and *PROTOCHLOROPHYLLIDE OXIDOREDUCTASE B* (*PORB*) transcripts are induced by *AtKAI2* signalling (**Figure 2D**). In this context, ethylene is a positive regulator of germination in a variety of species that is thought to inhibit ABA action.^25^ Ethylene also circumvents the requirement of SL for *Striga* germination.^26,27^ Similarly, activation of the KAI2 pathway leads to the induction of ethylene biosynthesis in *Lotus* roots.^28^ In summary, it appeared that activating AtKAI2 signalling increased GA and ethylene signalling and decreased ABA signalling. This combination of hormone action, which was necessary for germination in the dark, is also commonly seen during the germination of most plants in the light.^29^ This conserved pattern suggested that activation of KAI2/HTL signalling can substitute for light to dial up the correct germination code.

*Arabidopsis* thrives over a wide number of geographical ranges^30^, so locally adapted variation in *AtKAI2* signaling may contribute to the germination behaviour of some *Arabidopsis* populations. Because natural variation influences the *Arabidopsis* germination response to KARs in the light^31^, we decided to screen the ability of *Arabidopsis* seed collections from different geographical areas (accessions) to germinate in the dark. Thirty-seven accessions showed varying degrees of dark germination indicating natural divergence of this trait (**Figure 3A**). Of the three top accessions (Do-0, Copac-1, and Ara-1), Ara-1 showed elevated *AtKAI2* expression compared to the Col-0 accession (**Figure 3B and 3C**). To this point, Ara-1 seed mimicked *smax1-2* by germinating under levels of soil that inhibited Col-0 germination (**Figure 3D**). Moreover, transcript profiles of dark germinating Ara-1 seeds and *AtKAI2OX* Col-0 seed treated with KAR_2_ were highly correlated (r^2^=0.86), suggesting that these genotypes used similar genetic programs to germinate in the dark (**Figure 3E**). The increased expression of *AtKAI2* in Ara-1 seeds, however, only partially explains its dark germination phenotype, as this accession does not require KARs for dark germination. It has been hypothesized that *Arabidopsis* makes an endogenous ligand dubbed AtKAI2 ligand (KL), that activates AtKAI2.^32^ Natural variation in KL levels in *Arabidopsis* accessions could contribute to a spectrum of dark germination behaviours but this variation would have to be coupled with increased *AtKAI2* expression.

**Figure 3.**
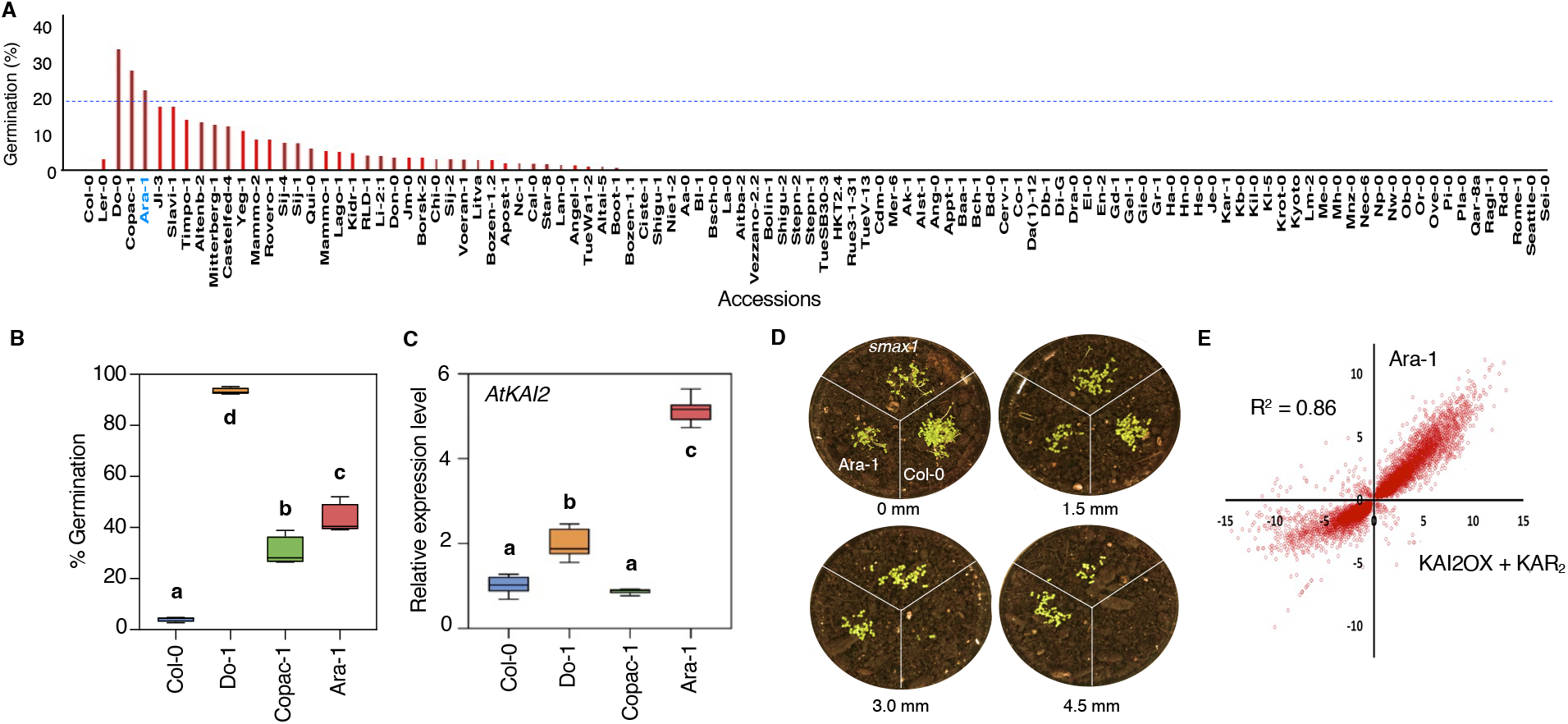
An *A. thaliana* accession, Ara-1 germinates in the dark and misexpresses *AtKAI2*. **(A)** The germination frequency of a collection of *Arabidopsis* accessions under dark conditions. The Ara-1 accession is highlighted in blue. The list of accessions is provided in Table S3. (**B**) Germination frequency of Col-0 and the top three dark germination lines (Ara-1, Do-1 and Copac-1) in the next generation. Each box represents three sets of biological replicates. (**C**) *AtKAI2* receptor expression of Col-0, Ara-1, Do-1, and Copac-1. RT–qPCR analysis of *AtKAI2* expression level relative to its expression in Col-0 seeds imbibed in the dark. An 18S rRNA gene was used as an internal control. Letters indicate significant differences (*P* < 0.05, ANOVA with post-hoc Tukey HSD Test). *P* values, sample sizes and box plot elements are provided in **Table S1**. (**D**) Representative picture of the germination of Col-0, *smax1-2* and Ara-1 seed buried at different depths of soil (1.5 to 4.5 cm). (**E**) Whole transcriptome comparison of 24h dark-germinating Ara-1 versus KAR_2_-treated *KAI2OX* seed. Log_2_-transformed values for probe responses for Ara-1 and *KAI2OX* + KAR_2_ are plotted on the *y-* and *x*-axis, respectively. The Pearson correlation coefficient (*r*^*2*^ = 0.86) is shown (degrees of freedom = 1, P-Value = 0, 99% confidence interval = 1.597281-1.647912). Three sets of independent biological replicates were used for transcriptome studies.

Interestingly, although dark-germinated Ara-1 and activated *AtKAI2OX* lines had very similar transcript profiles, expression of the *AtKAI2* homolog, *DWARF14-LIKE2* (*DLK2*) was reduced in Ara-1 but was highly induced in KAR_2_-treated *AtKAI2OX* seed (**Figure S2A**). The function of DLK2 is unknown but it does metabolize 2’*S-GR24*^33^ suggesting it may have roles in KAI2 signalling homeostasis. *DLK2* loss-of-function seeds did not germinate in the dark but they were sensitized to germinate under low light compared to wild type seed (**Figure S2B**). Furthermore, *kai2* mutants were epistatic to *dlk2* indicating *DLK2* functions genetically at or upstream of the receptor (**Figure S2B**). If DLK2 were involved in metabolizing KL, then loss of DLK2 function may result in higher levels of KL. Whatever the case, altered expression of both *AtKAI2* and *DLK2* in Ara-1 seeds suggested that this dark germinating accession, Ara-1 has aberrant AtKAI2 signalling compared to Col-0.

An *Arabidopsis KAI2*-dependent transcriptional signature for germination in the dark provides a window for looking at the germination code of experimentally intractable specialist plants like *Striga*. The ability of *AtKAI2* signalling to bypass light in *Arabidopsis* germination predicted that germinating *Striga* seed may show similar profiles. Indeed, many *Striga* gene homologs of *Arabidopsis* light-, GA-, ABA- and ethylene-responsive genes showed similar patterns of gene expression in germinating *Striga* seeds (**Figure 4**). Like KAR_2_-treated *AtKAI2OX* seed, SL stimulated expression of light-induced genes like LHCP, Rubisco, and components of Photosystems I and II in dark germinated *Striga* seed (**Figure 4**). In addition, the ability of SL to repress expression of ABA-induced genes is consistent with the observation that *Striga* seeds are capable of germinating on high concentrations of ABA.^34^

**Figure 4.**
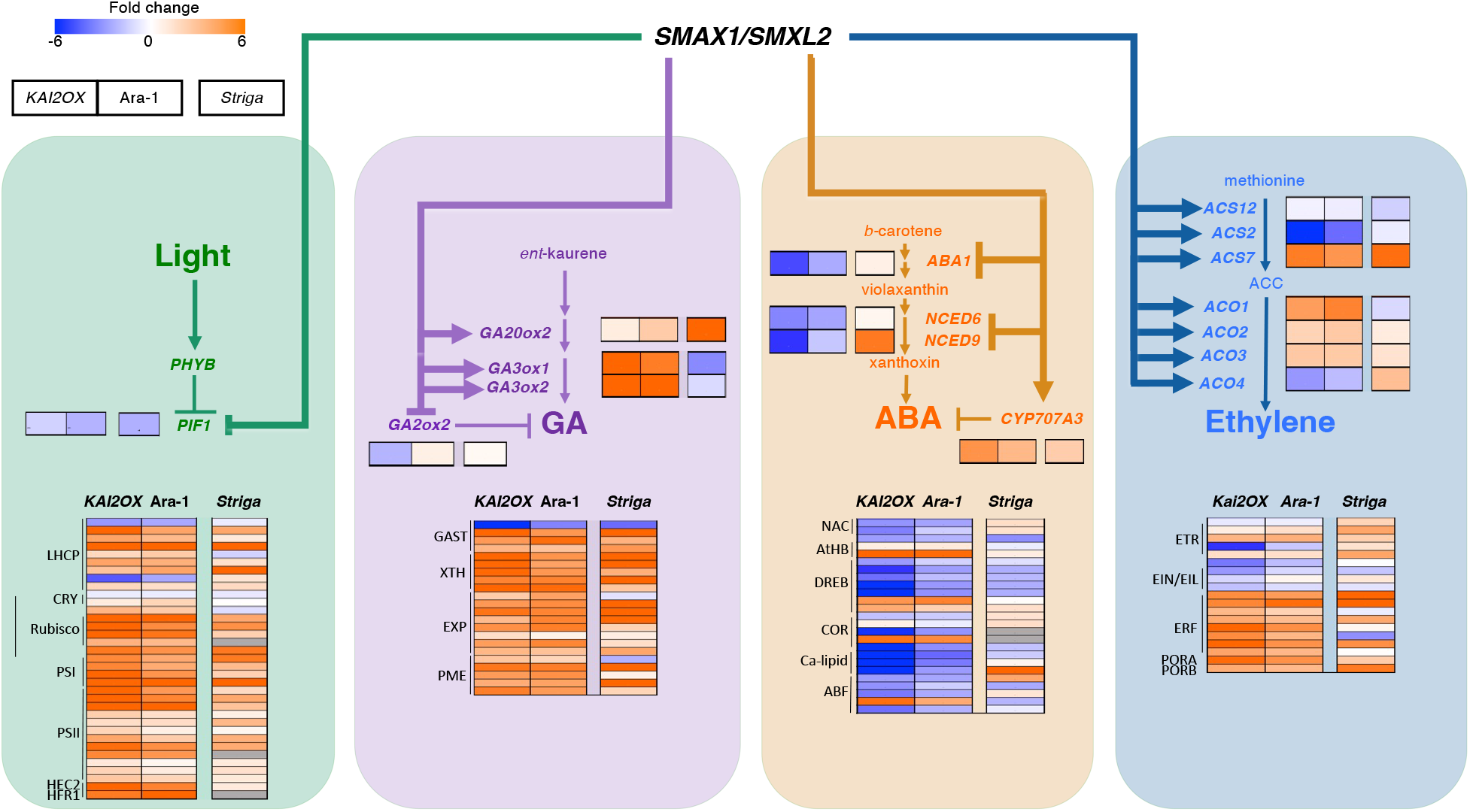
Expression of light and hormone regulated genes in germinating KAR_2_-treated *KAI2OX*, Ara-1, and *Striga* seeds. Heatmaps depict Log_2_ fold-change in gene expression during germination (24h) of KAR_2_ -treated *KAI2OX* (left box), Ara-1 (middle box), and *Striga* (right box) seeds. Partial biosynthetic pathways are shown for each hormone. Hormone-responsive genes are represented in panels. See **Table S2** for gene lists and values.

Interestingly, the expression of *Arabidopsis* and *Striga* genes involved in GA and ABA synthesis did not strongly correlate although we know levels of these hormones change in response to *rac*-GR24-treated *Striga* seed (**Figure 4**).^35^ This suggested that SL regulates GA and ABA levels differently in *Striga* versus *Arabidopsis*. By contrast, 1-AMINOCYCLOPROPANE-1-CARBOXYLATE SYNTHASE 7 (ACS7), a key enzyme in ethylene synthesis that is *AtKAI2*-inducible^28^ is upregulated in both species (**Figure 4A**). In summary, *HTL/KAI2* signalling dials up a correct code of light, GA, ABA, and ethylene responses that favour germination. This germination code appears to be conserved between a generalist plant like *Arabidopsis* which normally uses light as a germination cue and a specialist species like *Striga* which uses a small molecule cue.

Our analysis of HTL/KAI2 signalling pathway gives some insight into the evolution of germination signalling pathways in *Striga*. Adapting a signalling pathway for a particular process broadly follows scenarios where either the pathway is established for its function by natural selection (historical genesis) or the pathway, which has other functions, is co-opted for new uses (current utility).^36^ Although *HTL/KAI2* signalling has been implicated in a variety of developmental responses, our findings suggest its role in *Striga* germination is the product historical genesis rather than current utility. In generalist plants like *Arabidopsis*, SMAX1/SMXL2 inhibits germination in the absence of light similar to the repressive role of PIF1. Unlike PIF1, however, which is modulated by a ubiquitous environmental cue, light, SMAX1/SMXL2 levels are dependent on specific small molecules generated by distinct environmental conditions.^3,5^ Because SMAX1/SMXL2 function is necessary to repress dark germination, any selective pressures affecting their levels will dominate germination behaviours. We know substituting ShHTL receptors for AtKAI2 changes the small molecules that activate signalling and the magnitude of signal through the pathway.^10^ We also know only a few mutations are required to change a KAR receptor into an SL receptor.^37^ It is therefore easy to envision how selective pressures on HTL/KAI2 receptors contributed to the evolution of a conserved germination-signalling pathway. Our results emphasize that defining signalling pathways by their ligands and receptors can be confounding when these signals influence conserved downstream effectors that are also modulated by other signalling pathways. We suggest when looking at fundamental processes such as germination it is more informative to reference pathways by their bottom effectors like SMAX1.^38^

## Materials and Methods

### Plant Materials and Growth Conditions

All *Arabidopsis* strains, except the accession collection, are in the Col-0 background. We have previously described the generation of *KAI2OX, ShHTL7OX*, and *htl-3 dlk2-1* strains.^1,2^ The *smax1-2* mutant and *smax1-2 smxl2-1* double mutant were gifts from Dr. David Nelson.^3,4^ The *Arabidopsis* accessions from the 1001 genomes project were a gift from Dr. David Guttman. The *pif1-1* and *ein2-1* mutants were obtained from the Arabidopsis Biological Resource Center. *Striga hermonthica* seeds used in this study were collected from infested maize from the Kibos region in Kenya. All genotypes used in this study are described in the **Table S3**. All plants were grown under 24h, continuous white light at room temperature. Seeds were allowed to ripen for at least six months before use.

### Germination Assays

Our germination assays were performed on 0.8% agar plates supplemented with 10 μM benomyl. Stocks of GR24, KAR2, and GA_3_, were prepared in DMSO; paclobutrazol (PAC) was prepared in EtOH; and ABA was prepared in MeOH. For each experiment, we prepared a working stock 1,000 times the experimental concentration. Each stock was diluted 1:1,000 in 0.8% agar to a final solvent percentage of 0.1 (v/v). For all germination assays, the seeds were at least 6 months old. We emphasize that seeds ripened for less than 6 months have elevated background germination. For dark germination experiments, the seeds were sprinkled onto plates within 1 minute and then immediately wrapped in 3 layers of aluminum foil. For low light experiments, seeds were sprinkled onto plates and exposed to (0.3 µE) for specified durations, then wrapped in three layers of aluminum foil. Seeds were then stratified for 4 d at 4 °C and then placed in a dark cabinet at room temperature. After 5 days, germination was scored using radicle emergence. Dose response curves and effective concentration (EC_50_) values were generated with SigmaPlot 11.0.

### Accession Screen and Soil Germination Experiment

For germination experiments on agar, 104 accessions from the 1001 genome project were individually plated on 0.8% agar water medium supplemented with 10 μM benomyl. The plates were wrapped in three layers of aluminum foil within 1 minute of plating. After the seeds were plated, they were stratified for 4 d at 4°C and then placed in a dark cabinet at room temperature. After 5 days, germination was scored using radicle emergence.

For germination at different levels of soil, 0.6 g of damp soil was added to a 90 mm Petri dish. Approximately 10 mg of dry seeds per genotype were sprinkled on top. Damp soil was then sprinkled on top of the seeds to various levels. Plates were wrapped in three layers of aluminum foil and stratified at 4°C in the dark for 4 days. After this, the foil was removed from the top the plates and each plate was placed at room temperature in the light (15 µE) for 5 days.

### Double mutant construction

Putative *pif1 htl-3* double mutants were identified from F_2_ offspring of the cross between *pif1* and *htl-3; ShHTL7OX* by first screening for the *pif1* dark germination phenotype. Plants were later screened for the *htl-3* round rosette leaves and long hypocotyl phenotype. Potential *pif1 htl-3* homozygous double mutants were confirmed by genotyping of individual plants by PCR (see below).

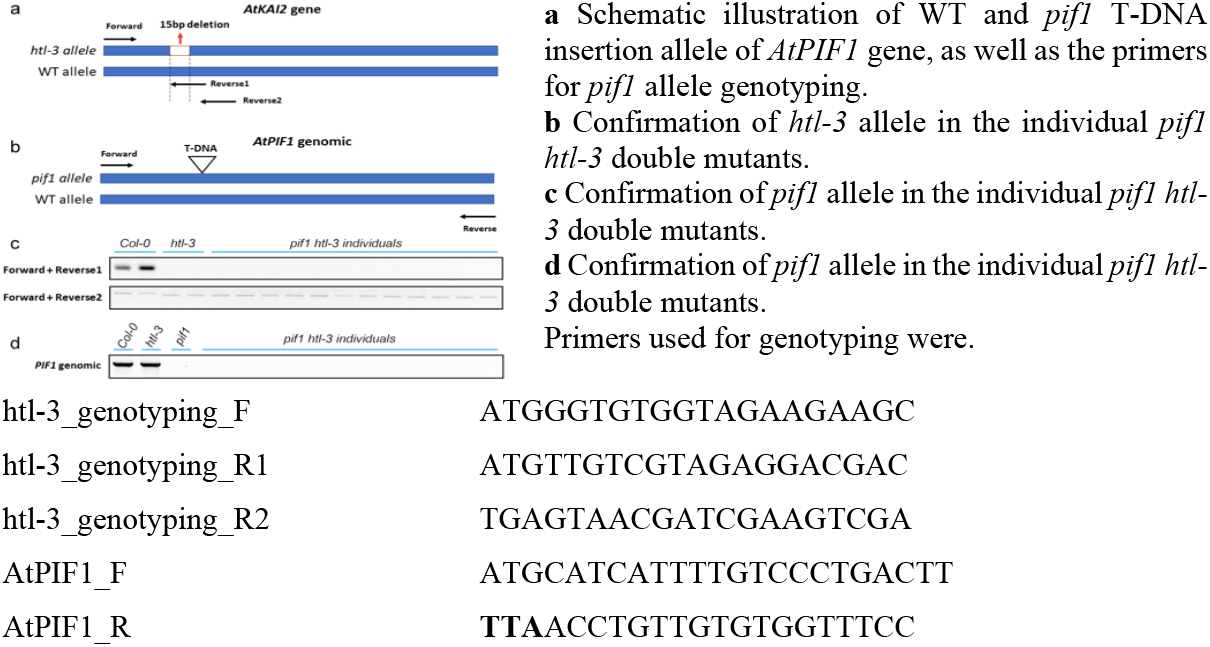

### Quantitative Gene Expression Analysis by RT-qPCR

#### Tissue Preparation and RNA Extraction

For *DLK2, KUF1*, and *BBX20* expression, seeds were sprinkled on 0.8% agar plates supplemented with vehicle or 0.5 μM KAR2 and immediately wrapped in aluminum foil. For *KAI2* expression, seeds were plated on 0.8% agar plates and immediately wrapped in foil. The seeds were stratified for 4 d at 4 °C and then placed in a dark cabinet at 25–26 °C. After 24 h, the seeds were flash-frozen in liquid N_2_. Seeds were then ground to a fine powder form using a frozen mortar and pestle. From this powder, total RNA was extracted in hot phenol (80 °C) using the phenol-chloroform extraction protocol, followed by treatment with DNase I.^5^ The samples were then purified with the Monarch® RNA Cleanup Kit (New England Biolabs).

#### Reverse transcription and qPCR

One μg of RNA was reverse transcribed using the LunaScript RT SuperMix Kit (New England Biolabs) according to the manual. One μl of 10-fold diluted cDNA was used in the qPCR. Primers (Supplementary Dataset) were designed using Beacon Designer 8 (PREMIER Biosoft International). Real-time PCR was performed using Luna Universal qPCR Master Mix (New England Biolabs) and quantified on a Bio-rad CFX96 Real-time Detection System (Bio-rad). Each reaction was performed in triplicate for each biological replicate. Expression levels were normalized against the expression of endogenous control gene, *AtACTIN8*, and were calculated using 2^-ΔΔCt^ method.^6^ All the expression data are presented as fold change value.

### Transcriptome studies

#### Tissue preparation and RNA extraction from Arabidopsis thaliana

*AtKAI2OX* seeds were plated on 0.8% agar plates supplemented with either 0.5 μM KAR2 or the vehicle; Ara-1 seeds were plated on a 0.8% agar plate supplemented with the vehicle. Each condition was prepared in triplicate. The plates were then immediately wrapped in aluminum foil and stratified for 4 d at 4 °C and then placed in a dark cabinet at room temperature. After 24h, the seeds were flash-frozen in liquid N_2_. Seeds were then ground to a fine powder form using a frozen mortar and pestle. From this powder, total RNA was extracted in hot phenol (80 °C) using the phenol-chloroform extraction protocol.

#### Transcript profiling and data analysis from Arabidopsis

The triplicate total RNA samples used for single-end 1×75bp RNA-Seq on a NextSeq500 sequencing platform (Illumina). After examining the data quality using FastQC (v0.11.5), the fastq data containing the raw reads for each sample was trimmed and filtered using htStream (v1.3.0) (UC, Davis) to generate clean reads. For each read, trimming and filtering processes include removing the last base, removing the low-quality fragments, removing ‘N’s and returning the longest fragments, and discarding reads that are less than 25 bp. Clean reads from each sample were subjected to HISAT2^**7**^ (v2.1.0) for genome mapping using *Arabidopsis thaliana* TAIR10 genome as reference. Samtools (v1.10) was used to remove the unmapped and multi-loci mapped reads and covert the uniquely mapped reads into a BAM file. Transcript assembly for each sample was performed using StringTie^**8**^ (v2.1.3) against *Arabidopsis thaliana* Araport11 annotation^**9**^ and transcriptome assemblies from each sample were merged using the *merge* function in StringTie to generate a combined transcriptome across all sequencing samples to serve as a new reference transcriptome. A customized python script, *stringtieGeneIdReplace2*.*py* (https://github.com/maplexuci/stringtie_gene_id_replacement) was written to replace the StringTie default ‘MSTRG’-tagged *gene_id* with the TAIR10 gene locus name to facilitate the comparisons of genes and transcripts between different samples. A StringTie provided python script, prepDE.py, was used to extract the read counts matrix from each transcriptome assemble and this counts matrix served as the input data for DESeq2 (v1.18.1).^**10**^ The data are found and described in NCBI GEO under GSE161704 entitled, *Natural variation in activation of KAI2/HTL pathway promotes germination in the dark in Arabidopsis*.

#### Tissue preparation and RNA extraction from Striga hermonthica

Forty mg aliquots of S. *hermonthica* seeds (collected from plants parasitizing maize in Kibos, Kenya in 2013) were surfaced sterilised with 10 % bleach for 7 min. Seeds were washed thoroughly with distilled water, placed on moistened filter paper (GF/A, Whatman™, Buckinghamshire) in Petri dishes and incubated in the dark at 30°C for 12 days. Seeds were collected at 12 days (prior to the addition of *rac* GR24) and 3 and 16 h after application of GR24. Seeds were washed from the plates onto clean GF/A filter paper. They were then placed in Eppendorf tubes and frozen in liquid nitrogen.

Total RNA was extracted using a RNeasy plant mini kit (Qiagen). Striga seeds were initially ground to a powder by placing two sterile 5 mm tungsten ball bearings into the Eppendorf tubes and shaking at 25 hz for 45s followed by an additional 60s, using a tissue lyser (Qiagen). Samples were eluted in 30 µl of dH_2_O. RNA was then extracted following the RNeasy plant mini kit protocol. RNA was eluted in 30 µl dH_2_O and the concentration determined using a Nanodrop-1000. RNA samples (3 µg) were treated with dsDNase (Thermocientific) to remove genomic DNA contamination following the manufacturer’s instructions. Samples were then purified and concentrated using a Monarch RNA Cleanup Kit (NEB). RNA samples were quantified and sent to BGI (China) for processing and sequencing.

#### Transcript profiling and data analysis from Striga hermonthica

Sequencing was performed on a BGISEQ-500 (BGI). Data quality was examined using FastQC (v0.11.5). Clean reads were obtained by trimming adaptors and filtering for quality score. After filtering, reads were mapped to *S. hermonthica* genome provided by J. Scholes^11^ using HISAT2. Gene expression levels were calculated using RSEM. Gene expression levels were calculated in Fragments Per Kilobase of transcript per Million mapped reads (FPKM). Sequencing data were deposited in NCBI under PRJNA802803, https://dataview.ncbi.nlm.nih.gov/object/PRJNA802803?reviewer=h9qigloj2glj899mome2m5mpla

Corresponding *A. thaliana* genes were identified in *S. hermonthica* by performing a BLAST analysis against the *S. hermonthica genome* provided by J. Scholes.^11^ *S. hermonthica* hits were confirmed by reciprocal BLAST to *A. thaliana* datasets. An e-value cut-off of 1e-20 was applied.

## Acknowledgements

We thank Dr. David Nelson for the gift of *smax1-2* and *smax1-2; smxl2-1*. We acknowledge the contribution of Dr. Peter McCourt for discussion and feedback on our manuscript. We acknowledge resource centers like the Arabidopsis Biological Resource Center (ABRC) for providing *Arabidopsis* mutants and accessions as well as the Parasitic Plant Genome Project for publicly available *Striga* transcriptome datasets. This work was supported by the New Frontiers in Research Fund Exploration program (2018-00118) and the Natural Sciences and Engineering Research Council of Canada (NSERC) in the form of a Discovery Grant (06752), an Accelerator Supplement (507992) and a Research Tools and Instruments grant awarded to S. Lumba.

## Author Contributions

S.L. contributed to all aspects of the research. M.B. designed and performed genetic and germination experiments, as well as analysed and interpreted data. Z.X. performed RNA-seq and qRT-PCR experiments on *Arabidopsis* and contributed to physiological assays. G.P. conducted germination assays of various genotypes. G.L. analysed RNA-seq data from *Arabidopsis* and *S. hermonthica*, identified homologs of *Arabidopsis* genes in *Striga hermonthica* and compared gene expression in both plants. J.H. performed germination assays. J.D.S. conducted RNA-seq experiments on *Striga*. C.S.P.M. and F-D.B. contributed compounds to conduct physiological experiments. S.L. and M.B. wrote the manuscript. S.L. directed and supervised all the research.

**Figure S1.**
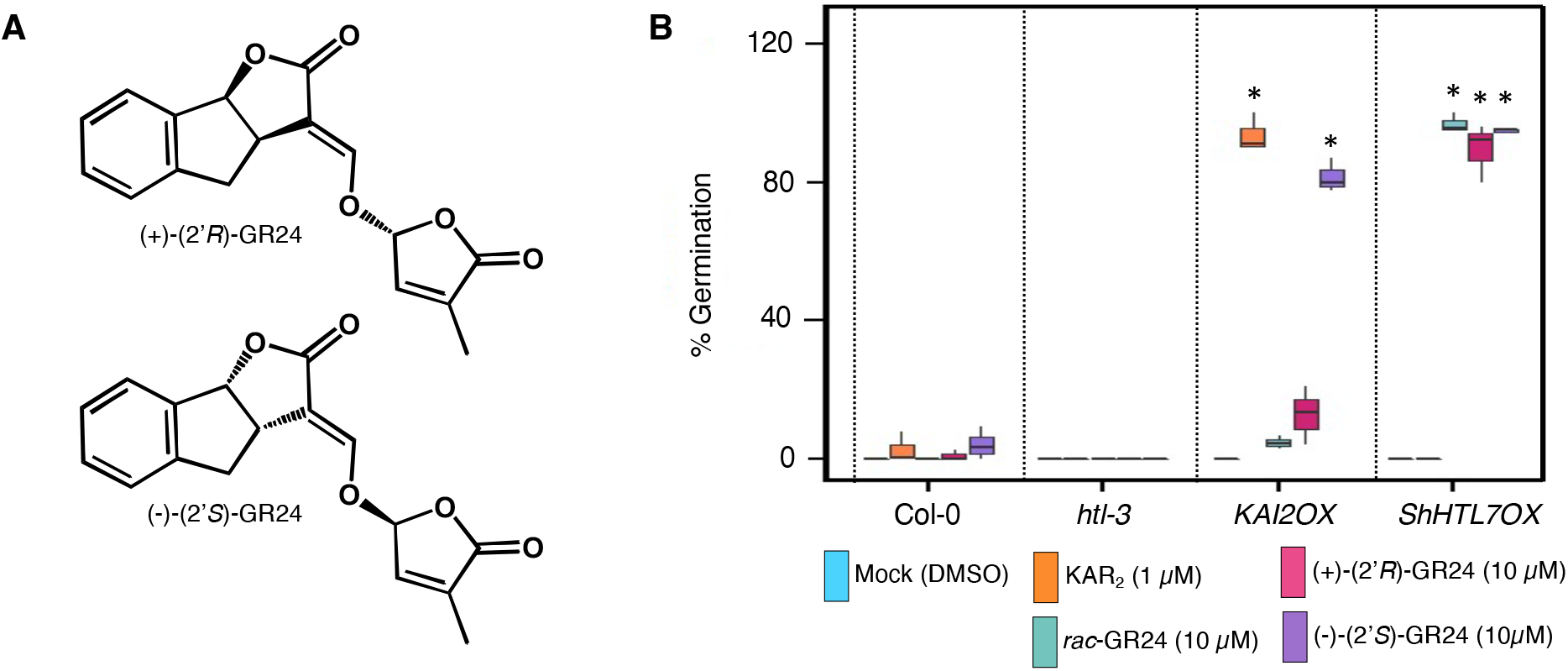
**A.** GR24 enantiomeric structures. **B.** Germination frequency of Col-0 (*wild type*) *htl-3* (loss-of-function allele of *AtKAI2*), *AtKAI2OX* (*35S::AtKAI2*) and *ShHTL7OX* ((*35S::ShHTL7*) seed on different GR24 enantiomers. Asterisk indicates a significant difference from the mock treatment (*P* < 0.01, ANOVA with Fisher’s LSD Test).

**Figure S2.**
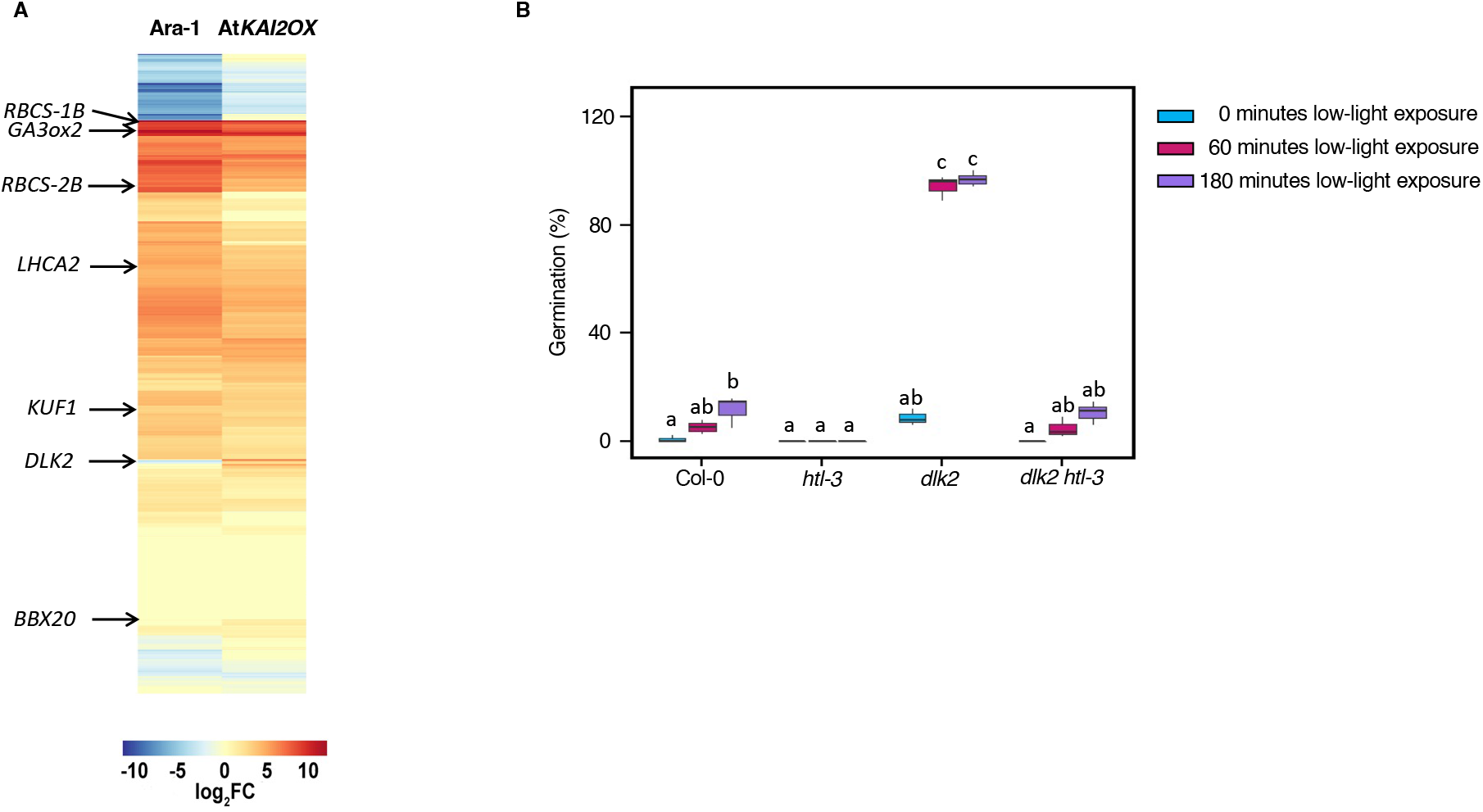
**A.** Heatmap representation of a subset of SL-regulated genes ^5^ where a gene is present in either the Ara-1 or AtKAI2OX+KAR_2_ dataset. This results in a 132 gene list which were then clustered. Genes of interest are shown with arrows. **B.** Germination frequency of Col-0 (*wild type*) *htl-3* (loss-of-function allele of *AtKAI2*), *dlk2* (loss-of-function allele of *DLK2*) and *dlk2 htl-3* double mutant seed germinated in the dark or upon exposure to low light (15*μ*E) for different time periods. Letters represent significant differences (P < 0.01, ANOVA with Fisher’s LSD Test).

