## Supplemental Table 1 for "HTL/KAI2 signalling substitutes for light to control plant germination"

**Table S1. Supplemental Table of Statistical Values**

**Fig. 1A**

**Sample distribution including sample size (n), mean, minima, median, maxima, 1st quartile, and 3rd quartile for Fig. 1A**

| Genotype | *Col-0* | | | | | | | | | |
|---|---|---|---|---|---|---|---|---|---|---|
| Condition | 0 minutes | 10 minutes | 20 minutes | 30 minutes | 40 minutes | 50 minutes | 60 minutes | 70 minutes | 120 minutes | 180 minutes |
| Sample Size | 3 | 3 | 3 | 3 | 3 | 3 | 3 | 3 | 3 | 3 |
| Min. | 0 | 0 | 0.000 | 0.800 | 2.247 | 1.282 | 0.000 | 6.173 | 12.28 | 41.56 |
| 1st Qu. | 0 | 0 | 0.000 | 2.839 | 2.552 | 2.070 | 2.679 | 7.574 | 13.90 | 41.87 |
| Median | 0 | 0 | 0.000 | 4.878 | 2.857 | 2.857 | 5.357 | 8.974 | 15.52 | 42.19 |
| Mean | 0 | 0 | 2.283 | 4.923 | 2.967 | 2.967 | 5.840 | 8.431 | 15.68 | 42.73 |
| 3rd Qu. | 0 | 0 | 3.425 | 6.984 | 3.327 | 3.810 | 8.760 | 9.560 | 17.37 | 43.32 |
| Max. | 0 | 0 | 6.849 | 9.091 | 3.797 | 4.762 | 12.162 | 10.145 | 19.23 | 44.44 |

| Genotype | *htl-3* | | | | | | | | | |
|---|---|---|---|---|---|---|---|---|---|---|
| Condition | 0 minutes | 10 minutes | 20 minutes | 30 minutes | 40 minutes | 50 minutes | 60 minutes | 70 minutes | 120 minutes | 180 minutes |
| Sample Size | 3 | 3 | 3 | 3 | 3 | 3 | 3 | 3 | 3 | 3 |
| Min. | 0 | 0 | 0 | 0 | 0.0000 | 0.000 | 0.000 | 1.639 | 2.632 | 4.878 |
| 1st Qu. | 0 | 0 | 0 | 0 | 0.0000 | 1.282 | 0.000 | 2.269 | 2.808 | 7.369 |
| Median | 0 | 0 | 0 | 0 | 0.0000 | 2.564 | 0.000 | 2.899 | 2.985 | 9.859 |
| Mean | 0 | 0 | 0 | 0 | 0.5952 | 4.141 | 1.709 | 2.612 | 4.539 | 10.041 |
| 3rd Qu. | 0 | 0 | 0 | 0 | 0.8929 | 6.212 | 2.564 | 3.098 | 5.493 | 12.622 |
| Max. | 0 | 0 | 0 | 0 | 1.7857 | 9.859 | 5.128 | 3.297 | 8.000 | 15.385 |

**Fig. 1C**

**Sample distribution including sample size (n), mean, minima, median, maxima, 1st quartile, and 3rd quartile for Fig. 1C**

| Genotype | *Col-0* | | | | | | | | | |
|---|---|---|---|---|---|---|---|---|---|---|
| Concentration of GR24 (µM) | 0 | 0.001 | 0.005 | 0.01 | 0.05 | 0.1 | 0.5 | 1 | 5 | 10 |
| Sample Size | 3 | 3 | 3 | 3 | 3 | 3 | 3 | 3 | 3 | 3 |
| Min. | 2.500 | 0 | 0.000 | 0 | 0 | 0.0000 | 0 | 0 | 0 | 0.0000 |
| 1st Qu. | 2.500 | 0 | 1.282 | 0 | 0 | 0.0000 | 0 | 0 | 0 | 0.0000 |
| Median | 2.500 | 0 | 2.564 | 0 | 0 | 0.0000 | 0 | 0 | 0 | 0.0000 |
| Mean | 2.816 | 0 | 2.336 | 0 | 0 | 0.7246 | 0 | 0 | 0 | 0.6667 |
| 3rd Qu. | 2.974 | 0 | 3.504 | 0 | 0 | 1.0870 | 0 | 0 | 0 | 1.0000 |
| Max. | 3.448 | 0 | 4.444 | 0 | 0 | 2.1739 | 0 | 0 | 0 | 2.0000 |

| Genotype | *htl-3* | | | | | | | | | |
|---|---|---|---|---|---|---|---|---|---|---|
| Concentration of GR24 (µM) | 0 | 0.001 | 0.005 | 0.01 | 0.05 | 0.1 | 0.5 | 1 | 5 | 10 |
| Sample Size | 3 | 3 | 3 | 3 | 3 | 3 | 3 | 3 | 3 | 3 |
| Min. | 0 | 0 | 0 | 0 | 0 | 0 | 0 | 0 | 0 | 0 |
| 1st Qu. | 0 | 0 | 0 | 0 | 0 | 0 | 0 | 0 | 0 | 0 |
| Median | 0 | 0 | 0 | 0 | 0 | 0 | 0 | 0 | 0 | 0 |
| Mean | 0 | 0 | 0 | 0 | 0 | 0 | 0 | 0 | 0 | 0 |
| 3rd Qu. | 0 | 0 | 0 | 0 | 0 | 0 | 0 | 0 | 0 | 0 |
| Max. | 0 | 0 | 0 | 0 | 0 | 0 | 0 | 0 | 0 | 0 |

| Genotype | *KAI2OX* | | | | | | | | | |
|---|---|---|---|---|---|---|---|---|---|---|
| Concentration of GR24 (µM) | 0 | 0.001 | 0.005 | 0.01 | 0.05 | 0.1 | 0.5 | 1 | 5 | 10 |
| Sample Size | 3 | 3 | 3 | 3 | 3 | 3 | 3 | 3 | 3 | 3 |
| Min. | 0.0000 | 0.0000 | 0 | 0 | 0 | 0 | 0.0000 | 0 | 0.0000 | 1.754 |
| 1st Qu. | 0.0000 | 0.0000 | 0 | 0 | 0 | 0 | 0.0000 | 0 | 0.8333 | 1.803 |
| Median | 0.0000 | 0.0000 | 0 | 0 | 0 | 0 | 0.0000 | 0 | 1.6667 | 1.852 |
| Mean | 0.5376 | 0.6944 | 0 | 0 | 0 | 0 | 0.7937 | 0 | 1.2222 | 1.831 |
| 3rd Qu. | 0.8065 | 1.0417 | 0 | 0 | 0 | 0 | 1.1905 | 0 | 1.8333 | 1.869 |
| Max. | 1.6129 | 2.0833 | 0 | 0 | 0 | 0 | 2.3810 | 0 | 2.0000 | 1.887 |

| Genotype | *ShHTL7* | | | | | | | | | |
|---|---|---|---|---|---|---|---|---|---|---|
| Concentration of GR24 (µM) | 0 | 0.001 | 0.005 | 0.01 | 0.05 | 0.1 | 0.5 | 1 | 5 | 10 |
| Sample Size | 3 | 3 | 3 | 3 | 3 | 3 | 3 | 3 | 3 | 3 |
| Min. | 0 | 0.000 | 13.64 | 19.61 | 38.33 | 46.43 | 61.19 | 64.58 | 70.00 | 70.27 |
| 1st Qu. | 0 | 0.000 | 14.77 | 20.15 | 42.75 | 47.41 | 67.55 | 67.29 | 73.24 | 73.07 |
| Median | 0 | 0.000 | 15.91 | 20.69 | 47.17 | 48.39 | 73.91 | 70.00 | 76.47 | 75.86 |
| Mean | 0 | 1.905 | 15.70 | 22.79 | 47.80 | 50.00 | 70.59 | 69.86 | 75.65 | 78.94 |
| 3rd Qu. | 0 | 2.857 | 16.73 | 24.38 | 52.53 | 51.78 | 75.29 | 72.50 | 78.48 | 83.28 |
| Max. | 0 | 5.714 | 17.54 | 28.07 | 57.89 | 55.17 | 76.67 | 75.00 | 80.49 | 90.70 |

**Fig. 1D**

**Sample distribution including sample size (n), mean, minima, median, maxima, 1st quartile, and 3rd quartile for Fig. 1D**

| Genotype | *Col-0* | | | | | | | | | |
|---|---|---|---|---|---|---|---|---|---|---|
| Concentration KAR2 (µM) | 0 | 0.001 | 0.005 | 0.01 | 0.05 | 0.1 | 0.5 | 1 | 5 | 10 |
| Sample Size | 3 | 3 | 3 | 3 | 3 | 3 | 3 | 3 | 3 | 3 |
| Min. | 0 | 0.000 | 0 | 0 | 0.0000 | 0.0000 | 0 | 0 | 0.0000 | 0 |
| 1st Qu. | 0 | 1.220 | 0 | 0 | 0.0000 | 0.0000 | 0 | 0 | 0.0000 | 0 |

| | | | | | | | | | | |
|---|---|---|---|---|---|---|---|---|---|---|
| Median | 0 | 2.439 | 0 | 0 | 0.0000 | 0.0000 | 0 | 0 | 0.0000 | 0 |
| Mean | 0 | 1.668 | 0 | 0 | 0.8439 | 0.9259 | 0 | 0 | 0.6667 | 0 |
| 3rd Qu. | 0 | 2.502 | 0 | 0 | 1.2658 | 1.3889 | 0 | 0 | 1.0000 | 0 |
| Max. | 0 | 2.564 | 0 | 0 | 2.5316 | 2.7778 | 0 | 0 | 2.0000 | 0 |

| Genotype | *ShHTL7* | | | | | | | | | |
|---|---|---|---|---|---|---|---|---|---|---|
| Concentration KAR2 (μM) | 0 | 0.001 | 0.005 | 0.01 | 0.05 | 0.1 | 0.5 | 1 | 5 | 10 |
| Sample Size | 3 | 3 | 3 | 3 | 3 | 3 | 3 | 3 | 3 | 3 |
| Min. | 0 | 0 | 2.500 | 0 | 0 | 0.0000 | 0 | 0 | 0.0000 | 0 |
| 1st Qu. | 0 | 0 | 2.917 | 0 | 0 | 0.6849 | 0 | 0 | 0.0000 | 0 |
| Median | 0 | 0 | 3.333 | 0 | 0 | 1.3699 | 0 | 0 | 0.0000 | 0 |
| Mean | 0 | 0 | 4.843 | 0 | 0 | 1.3825 | 0 | 0 | 0.6410 | 0 |
| 3rd Qu. | 0 | 0 | 6.014 | 0 | 0 | 2.0738 | 0 | 0 | 0.9615 | 0 |
| Max. | 0 | 0 | 8.696 | 0 | 0 | 2.7778 | 0 | 0 | 1.9231 | 0 |

| Genotype | *AtHTL* Replicate 1 | | | | | | | | | |
|---|---|---|---|---|---|---|---|---|---|---|
| Concentration KAR2 (μM) | 0 | 0.001 | 0.005 | 0.01 | 0.05 | 0.1 | 0.5 | 1 | 5 | 10 |
| Sample Size | 3 | 3 | 3 | 3 | 3 | 3 | 3 | 3 | 3 | 3 |
| Min. | 0.000 | 5.714 | 25.00 | 57.58 | 67.27 | 57.14 | 58.82 | 61.29 | 65.62 | 61.54 |
| 1st Qu. | 1.613 | 8.413 | 28.12 | 59.28 | 67.64 | 63.99 | 62.20 | 63.98 | 68.66 | 64.64 |
| Median | 3.226 | 11.111 | 31.25 | 60.98 | 68.00 | 70.83 | 65.57 | 66.67 | 71.70 | 67.74 |
| Mean | 2.830 | 11.749 | 34.47 | 62.43 | 68.90 | 66.57 | 63.44 | 68.09 | 69.77 | 67.17 |
| 3rd Qu. | 4.244 | 14.766 | 39.21 | 64.86 | 69.71 | 71.29 | 65.74 | 71.49 | 71.85 | 69.98 |
| Max. | 5.263 | 18.421 | 47.17 | 68.75 | 71.43 | 71.74 | 65.91 | 76.32 | 72.00 | 72.22 |

| Genotype | *AtHTL* Replicate 2 | | | | | | | | | |
|---|---|---|---|---|---|---|---|---|---|---|
| Concentration KAR2 (μM) | 0 | 0.001 | 0.005 | 0.01 | 0.05 | 0.1 | 0.5 | 1 | 5 | 10 |
| Sample Size | 3 | 3 | 3 | 3 | 3 | 3 | 3 | 3 | 3 | 3 |
| Min. | 3.125 | 17.86 | 41.67 | 62.96 | 65.38 | 73.08 | 73.81 | 76.92 | 62.50 | 64.71 |
| 1st Qu. | 3.381 | 18.93 | 44.36 | 67.20 | 70.19 | 76.54 | 73.86 | 77.94 | 72.32 | 71.75 |
| Median | 3.636 | 20.00 | 47.06 | 71.43 | 75.00 | 80.00 | 73.91 | 78.95 | 82.14 | 78.79 |
| Mean | 3.643 | 19.71 | 49.33 | 70.38 | 75.00 | 78.41 | 75.68 | 79.54 | 76.11 | 74.22 |
| 3rd Qu. | 3.902 | 20.64 | 53.16 | 74.09 | 79.81 | 81.07 | 76.61 | 80.85 | 82.91 | 78.98 |
| Max. | 4.167 | 21.28 | 59.26 | 76.74 | 84.62 | 82.14 | 79.31 | 82.76 | 83.67 | 79.17 |

**Fig. 1E**

**Box-plot elements including sample size (n), mean, minima, median, maxima, 1st quartile, 3rd quartile, p-values and p-value comparisons are listed for Figure 1E.**

*DLK2* Expression

| Genotype | Col-0 | *KAI2OX* | *htl-3* | *smax1-2* | *smax1-2; smxl2-1* | Col-0 | *KAI2OX* | *htl-3* | *smax1-2* | *smax1-2; smxl2-1* |
|---|---|---|---|---|---|---|---|---|---|---|
| Condition | Mock | Mock | Mock | Mock | Mock | 0.5 μM KAR2 | 0.5 μM KAR2 | 0.5 μM KAR2 | 0.5 μM KAR2 | 0.5 μM KAR2 |
| Sample Size | 9 | 9 | 9 | 9 | 9 | 9 | 9 | 9 | 9 | 9 |
| Min. | -0.29339 | -1.35341 | -6.389 | 0.2762 | 3.060 | -0.71921 | 2.709 | -5.957 | 0.9006 | 2.611 |
| 1st Qu. | -0.14477 | -1.00701 | -5.725 | 0.5967 | 3.173 | -0.38142 | 2.928 | -5.650 | 0.9812 | 3.202 |
| Median | 0.03747 | -0.74712 | -5.283 | 0.8077 | 3.250 | -0.27092 | 2.990 | -5.082 | 1.2825 | 3.561 |
| Mean | -0.01182 | -0.70384 | -5.470 | 0.6970 | 3.325 | -0.20599 | 2.957 | -5.135 | 1.2072 | 3.481 |
| 3rd Qu. | 0.13798 | -0.34002 | -5.204 | 0.8425 | 3.361 | 0.03177 | 3.053 | -4.937 | 1.3583 | 3.932 |
| Max. | 0.26458 | 0.01933 | -4.746 | 0.9480 | 3.968 | 0.25706 | 3.130 | -3.546 | 1.5963 | 4.254 |

*KUF1* Expression

| Genotype | Col-0 | *KAI2OX* | *htl-3* | *smax1-2* | *smax1-2; smxl2-1* | Col-0 | *KAI2OX* | *htl-3* | *smax1-2* | *smax1-2; smxl2-1* |
|---|---|---|---|---|---|---|---|---|---|---|
| Condition | Mock | Mock | Mock | Mock | Mock | 0.5 μM KAR2 | 0.5 μM KAR2 | 0.5 μM KAR2 | 0.5 μM KAR2 | 0.5 μM KAR2 |
| Sample Size | 9 | 9 | 9 | 9 | 9 | 9 | 9 | 9 | 9 | 9 |
| Min. | -0.72968 | -2.5880 | -2.870 | 2.541 | 3.645 | -0.3182 | 2.869 | -3.513 | 2.791 | 3.593 |
| 1st Qu. | -0.39137 | -1.4922 | -2.606 | 2.734 | 3.663 | 0.1609 | 3.137 | -2.850 | 2.881 | 3.880 |
| Median | 0.07822 | -1.2692 | -2.280 | 2.847 | 4.082 | 0.2557 | 3.354 | -2.551 | 2.929 | 4.317 |
| Mean | -0.05694 | -1.1831 | -2.307 | 2.845 | 4.101 | 0.3046 | 3.283 | -2.423 | 3.062 | 4.247 |
| 3rd Qu. | 0.15141 | -0.7362 | -2.003 | 2.992 | 4.270 | 0.5535 | 3.406 | -2.221 | 3.237 | 4.535 |
| Max. | 0.47879 | -0.1384 | -1.840 | 3.157 | 5.044 | 0.7821 | 3.662 | -1.250 | 3.527 | 4.868 |

*BBX20* Expression

| Genotype | Col-0 | *KAI2OX* | *htl-3* | *smax1-2* | *smax1-2; smxl2-1* | Col-0 | *KAI2OX* | *htl-3* | *smax1-2* | *smax1-2; smxl2-1* |
|---|---|---|---|---|---|---|---|---|---|---|
| Condition | Mock | Mock | Mock | Mock | Mock | 0.5 μM KAR2 | 0.5 μM KAR2 | 0.5 μM KAR2 | 0.5 μM KAR2 | 0.5 μM KAR2 |
| Sample Size | 9 | 9 | 9 | 9 | 9 | 9 | 9 | 9 | 9 | 9 |
| Min. | -0.45682 | -1.1856 | -4.854 | 0.1419 | 0.3786 | -0.8057 | 0.9198 | -4.426 | 0.07566 | 0.863 |
| 1st Qu. | -0.22036 | -1.0785 | -4.010 | 0.2523 | 0.8308 | -0.4494 | 1.0459 | -4.251 | 0.33075 | 1.310 |
| Median | 0.09910 | -1.0570 | -3.734 | 0.3995 | 1.2277 | -0.3189 | 1.1917 | -3.947 | 0.78977 | 1.716 |
| Mean | -0.02303 | -0.8301 | -3.883 | 0.4345 | 1.1985 | -0.3423 | 1.2553 | -3.958 | 0.65312 | 1.507 |
| 3rd Qu. | 0.19717 | -0.6087 | -3.711 | 0.5676 | 1.4593 | -0.1248 | 1.4893 | -3.770 | 0.88197 | 1.779 |
| Max. | 0.28169 | -0.2046 | -3.507 | 0.8542 | 1.9183 | -0.0417 | 1.8354 | -3.184 | 0.95452 | 1.867 |

**Fig. 2**

**Box-plot elements including sample size (n), mean, minima, median, maxima, 1st quartile, 3rd quartile, p-values are listed for Figure 2.**

| | Fig. 2B | | Fig. 2C | | Fig. 2D | |
|---|---|---|---|---|---|---|
| Genotype | *ShHTL7* | *ShHTL7* | *ShHTL7* | *ShHTL7* | *ShHTL7* | *ShHTL7; ein2-1* |
| Condition | 1 µM GR24 | 1 µM GR24 + 20 µM PAC | 1 µM GR24 | 1 µM GR24 + 3 µM ABA | 1 µM GR24 | 1 µM GR24 |
| Min. | 93.48 | 0 | 91.53 | 53.33 | 97.44 | 11.76 |
| 1st Qu. | 95.49 | 0 | 91.84 | 58.58 | 97.70 | 12.27 |
| Median | 97.50 | 0 | 92.16 | 63.83 | 97.96 | 12.77 |
| Mean | 96.33 | 0 | 92.71 | 62.74 | 98.47 | 13.53 |
| 3rd Qu. | 97.75 | 0 | 93.30 | 67.44 | 98.98 | 14.42 |
| Max. | 98.00 | 0 | 94.44 | 71.05 | 100.00 | 16.07 |
| P-value | 7.24E-10 | | 1.21E-03 | | 2.19E-09 | |

**Fig. 3B**

**Box-plot elements including sample size (n), mean, minima, median, maxima, 1st quartile, 3rd quartile, p-values and p-value comparisons are listed for Figure 3b.**

| | Fig. 3B | | | |
|---|---|---|---|---|
| Genotype | Col-0 | Do-0 | Copac-1 | Ara-1 |
| Sample Size | 4 | 4 | 4 | 4 |
| Min. | 2.740 | 92.31 | 26.47 | 39.13 |
| 1st Qu. | 3.947 | 92.59 | 26.81 | 39.78 |
| Median | 4.485 | 92.93 | 28.09 | 40.45 |
| Mean | 4.117 | 93.35 | 30.39 | 43.01 |
| 3rd Qu. | 4.655 | 93.69 | 31.68 | 43.68 |
| Max. | 4.760 | 95.24 | 38.89 | 52.00 |

***P*-values of one-way ANOVA with Tukey HSD multiple comparison for dark germination rate of *Col-0*, *Do-1*, *Copac-1* and *Ara-1***

| ***% Dark Germination*** | Col-0 | Do-0 | Copac-1 | Ara-1 |
|---|---|---|---|---|
| Col-0 | - | - | - | - |
| Do-0 | 8.46E-12 | | - | - |
| Capac-1 | 8.06E-06 | 4.48E-10 | | - |
| Ara-1 | 1.13E-07 | 6.01E-09 | 5.97E-03 | - |

**Fig. 3C**

**Box-plot elements including sample size (n), mean, minima, median, maxima, 1st quartile, 3rd quartile, p-values and p-value comparisons are listed for Figure 3C.**

| | Fig. 3C | | | |
|---|---|---|---|---|
| Genotype | Col-0 | Do-0 | Copac-1 | Ara-1 |
| Sample Size | 9 | 9 | 9 | 9 |
| Min. | -0.53565 | 0.6379 | -0.3826 | 2.242 |
| 1st Qu. | -0.18361 | 0.8078 | -0.1988 | 2.321 |
| Median | 0.03082 | 0.9091 | -0.1587 | 2.368 |
| Mean | 0.00000 | 0.9730 | -0.1936 | 2.358 |
| 3rd Qu. | 0.28004 | 1.2231 | -0.1227 | 2.390 |
| Max. | 0.35159 | 1.3004 | -0.1069 | 2.496 |

**Supplemental Figure 1**

**Box-plot elements including sample size (n), mean, minima, median, maxima, 1st quartile, 3rd quartile, p-values and p-value comparisons are listed for Supplementary Figure 1.**

| Genotype | *Col-0* | | | | | *htl-3* | | | | |
|---|---|---|---|---|---|---|---|---|---|---|
| Condition | Mock | 1 µM KAR2 | 10 µM rac-GR24 | 10 µM (+)-GR24 | 10 µM (-)-GR24 | Mock | 1 µM KAR2 | 10 µM rac-GR24 | 10 µM (+)-GR24 | 10 µM (-)-GR24 |
| Sample Size | 3 | 3 | 3 | 3 | 3 | 3 | 3 | 3 | 3 | 3 |
| Min. | 0 | 0.0000 | 0 | 0.0000 | 0.000 | 0 | 0 | 0 | 0 | 0 |
| 1st Qu. | 0 | 0.2857 | 0 | 0.0000 | 1.667 | 0 | 0 | 0 | 0 | 0 |
| Median | 0 | 0.5714 | 0 | 0.0000 | 3.333 | 0 | 0 | 0 | 0 | 0 |
| Mean | 0 | 2.8571 | 0 | 0.9259 | 4.286 | 0 | 0 | 0 | 0 | 0 |
| 3rd Qu. | 0 | 4.2857 | 0 | 1.3889 | 6.429 | 0 | 0 | 0 | 0 | 0 |
| Max. | 0 | 8.0000 | 0 | 2.7778 | 9.524 | 0 | 0 | 0 | 0 | 0 |
| p-value | N/A | 2.70E-01 | 1 | 7.15E-01 | 1.09E-01 | N/A | 1 | 1 | 1 | 1 |
| Signficant | | No | No | No | No | | No | No | No | No |

| Genotype | *AtHTL* | | | | | *ShHTL7* | | | | |
|---|---|---|---|---|---|---|---|---|---|---|
| Condition | Mock | 1 µM KAR2 | 10 µM rac-GR24 | 10 µM (+)-GR24 | 10 µM (-)-GR24 | Mock | 1 µM KAR2 | 10 µM rac-GR24 | 10 µM (+)-GR24 | 10 µM (-)-GR24 |
| Sample Size | 3 | 3 | 3 | 3 | 3 | 3 | 3 | 3 | 3 | 3 |
| Min. | 0 | 90.00 | 3.125 | 4.000 | 77.42 | 0 | 0 | 95.00 | 80.00 | 93.94 |
| 1st Qu. | 0 | 90.45 | 3.736 | 8.818 | 78.71 | 0 | 0 | 95.33 | 86.15 | 94.59 |
| Median | 0 | 90.91 | 4.348 | 13.636 | 80.00 | 0 | 0 | 95.65 | 92.31 | 95.24 |
| Mean | 0 | 93.64 | 4.713 | 12.823 | 81.46 | 0 | 0 | 96.88 | 89.44 | 94.88 |
| 3rd Qu. | 0 | 95.45 | 5.507 | 17.235 | 83.48 | 0 | 0 | 97.83 | 94.15 | 95.35 |
| Max. | 0 | 100.00 | 6.667 | 20.833 | 86.96 | 0 | 0 | 100.00 | 96.00 | 95.45 |

| | | | | | | | | | | |
|---|---|---|---|---|---|---|---|---|---|---|
| p-value | N/A | 3.45E-11 | 2.79E-01 | 9.43E-03 | 3.52E-10 | N/A | 1 | 1.18E-12 | 3.04E-12 | 1.51E-12 |
| Signficant | | Yes | No | No | Yes | | No | Yes | Yes | Yes |

**Supplemental Figure 2**

**Box-plot elements including sample size (n), mean, minima, median, maxima, 1st quartile, 3rd quartile, p-values and p-value comparisons are listed for Supplementary Figure 2.**

| Genotype | *Col-0* | | | *htl-3* | | |
|---|---|---|---|---|---|---|
| Light Exposure (Minutes) | 0 | 60 | 180 | 0 | 60 | 180 |
| Sample Size | 3 | 3 | 3 | 3 | 3 | 3 |
| Min. | 0.0000 | 2.564 | 4.878 | 0 | 0 | 0 |
| 1st Qu. | 0.0000 | 3.914 | 9.731 | 0 | 0 | 0 |
| Median | 0.0000 | 5.263 | 14.583 | 0 | 0 | 0 |
| Mean | 0.7092 | 5.276 | 11.695 | 0 | 0 | 0 |
| 3rd Qu. | 1.0638 | 6.632 | 15.104 | 0 | 0 | 0 |
| Max. | 2.1277 | 8.000 | 15.625 | 0 | 0 | 0 |

| Genotype | *dlk2* | | | *dlk2 htl* | | |
|---|---|---|---|---|---|---|
| Light Exposure (Minutes) | 0 | 60 | 180 | 0 | 60 | 180 |
| Sample Size | 3 | 3 | 3 | 3 | 3 | 3 |
| Min. | 6.061 | 88.89 | 93.94 | 0 | 2.000 | 6.061 |
| 1st Qu. | 6.978 | 92.44 | 95.27 | 0 | 2.667 | 8.586 |
| Median | 7.895 | 96.00 | 96.61 | 0 | 3.333 | 11.111 |
| Mean | 8.692 | 94.11 | 96.85 | 0 | 4.808 | 10.585 |
| 3rd Qu. | 10.008 | 96.72 | 98.31 | 0 | 6.212 | 12.847 |
| Max. | 12.121 | 97.44 | 100.00 | 0 | 9.091 | 14.583 |

***P*-values of one-way ANOVA with Fisher LSD multiple comparison**

| | **Col-0 (0 min.)** | **Col-0 (60 min.)** | **Col-0 (180 min.)** | **dlk2 (0 min.)** | **dlk2 (60 min.)** | **dlk2 (180 min.)** | **dlk2 htl (0 min.)** | **dlk2 htl (60 min.)** | **dlk2 htl (180 min.)** | **htl (0 min.)** | **htl (60 min.)** |
|---|---|---|---|---|---|---|---|---|---|---|---|
| **Col-0 (0 min.)** | - | - | - | - | - | - | - | - | - | - | - |
| **Col-0 (60 min.)** | 0.08419843 | - | - | - | - | - | - | - | - | - | - |
| **Col-0 (180 min.)** | 0.00020225 | 0.01803918 | - | - | - | - | - | - | - | - | - |
| **dlk2 (0 min.)** | 0.00419015 | 0.19077239 | 0.24833985 | - | - | - | - | - | - | - | - |
| **dlk2 (60 min.)** | 0 | 0 | 0 | 0 | - | - | - | - | - | - | - |
| **dlk2 (180 min.)** | 0 | 0 | 0 | 0 | 0.29093908 | - | - | - | - | - | - |
| **dlk2 htl (0 min.)** | 0.78255543 | 0.04807894 | 9.70E-05 | 0.00208909 | 0 | 0 | - | - | - | - | - |
| **dlk2 htl (60 min.)** | 0.1191097 | 0.85552801 | 0.01180497 | 0.13878077 | 0 | 0 | 0.06988246 | - | - | - | - |
| **dlk2 htl (180 min.)** | 0.00063362 | 0.04677785 | 0.66598846 | 0.46339466 | 0 | 0 | 0.00030607 | 0.03163027 | - | - | - |
| **htl (0 min.)** | 0.78255543 | 0.04807894 | 9.70E-05 | 0.00208909 | 0 | 0 | 1 | 0.06988246 | 0.00030607 | - | - |
| **htl (60 min.)** | 0.78255543 | 0.04807894 | 9.70E-05 | 0.00208909 | 0 | 0 | 1 | 0.06988246 | 0.00030607 | 1 | - |
| **htl (180 min.)** | 0.78255543 | 0.04807894 | 9.70E-05 | 0.00208909 | 0 | 0 | 1 | 0.06988246 | 0.00030607 | 1 | 1 |
