## Supplemental Table 3 for "HTL/KAI2 signalling substitutes for light to control plant germination"

**Table S3. Supplemental Table of Genotypes Used in this Study**

| Genotype | Background | ABRC Stock number |
|---|---|---|
| Col-0 | Col-0 | N/A |
| Ler-0 | Ler-0 | N/A |
| *htl-3/kai2-3* | Col-0 | N/A |
| *smax1-2* | Col-0 | SALK_128579 (Nelson lab) |
| *smax1-2 smxl2-1* | Col-0 | N/A |
| *htl-3 35S::AtKAI2* | Col-0 | N/A |
| *htl-3 35S::ShHTL7* | Col-0 | N/A |
| *htl-3 pif1-1* | Col-0 | N/A |
| *ein2-1 35S::ShHTL7* | Col-0 | N/A |
| Aa-0 | Aa-0 | CS76428 |
| Aitba-2 | Aitba-2 | CS76347 |
| Ak-1 | Ak-1 | CS76431 |
| Alst-1 | Alst-1 | CS76432 |
| Altai-5 | Altai-5 | CS76433 |
| Altai-5 | Altai-5 | CS76433 |
| Altenb-2 | Altenb-2 | CS76353 |
| Ang-0 | Ang-0 | CS76436 |
| Angel-1 | Angel-1 | CS76362 |
| Apost-1 | Apost-1 | CS76368 |
| Appt-1 | Appt-1 | CS76440 |
| Ara-1 | Ara-1 | CS76382 |
| Baa-1 | Baa-1 | CS76442 |
| Bch-1 | Bch-1 | CS76444 |
| Bd-0 | Bd-0 | CS76445 |
| Bl-1 | Bl-1 | CS76450 |
| Bolin-1 | Bolin-1 | CS76373 |
| Boot-1 | Boot-1 | CS76452 |
| Borsk-2 | Borsk-2 | CS76421 |
| Bozen-1.1 | Bozen-1.1 | CS76357 |
| Bozen-1.2 | Bozen-1.2 | CS76358 |
| Bsch-0 | Bsch-0 | CS76457 |
| Cal-0 | Cal-0 | CS76460 |
| Castelfed-4-212 | Castelfed-4-212 | CS76355 |
| Cdm-0 | Cdm-0 | CS76410 |
| Cerv-1 | Cerv-1 | CS76462 |
| Chi-0 | Chi-0 | CS76464 |
| Ciste-1 | Ciste-1 | CS76359 |
| Co-1 | Co-1 | CS76468 |
| Copac-1 | Copac-1 | CS76420 |
| Da(1)-12 | Da(1)-12 | CS76470 |
| Db-1 | Db-1 | CS76471 |
| Di-G | Di-G | CS76472 |
| Do-0 | Do-0 | CS76474 |
| Don-0 | Don-0 | CS76411 |
| Dra-0 | Dra-0 | CS76476 |
| El-0 | El-0 | CS76479 |
| En-2 | En-2 | CS76481 |
| Gd-1 | Gd-1 | CS76491 |
| Gel-1 | Gel-1 | CS76492 |
| Gie-0 | Gie-0 | CS76493 |
| Gr-1 | Gr-1 | CS76496 |
| Ha-0 | Ha-0 | CS76500 |
| HKT2.4 | HKT2.4 | CS76404 |
| Hn-0 | Hn-0 | CS76513 |
| Hs-0 | Hs-0 | CS76515 |
| Je-0 | Je-0 | CS76518 |
| Jl-3 | Jl-3 | CS76519 |
| Jm-0 | Jm-0 | CS76520 |
| Kar-1 | Kar-1 | CS76522 |
| Kb-0 | Kb-0 | CS76524 |
| Kidr-1 | Kidr-1 | CS76376 |
| Kil-0 | Kil-0 | CS76526 |
| Kl-5 | Kl-5 | CS76528 |
| Krot-0 | Krot-0 | CS76534 |
| Kyoto | Kyoto | CS76535 |
| La-0 | La-0 | CS76538 |
| Lago-1 | Lago-1 | CS76367 |
| Lan-0 | Lan-0 | CS76539 |
| Li-2:1 | Li-2:1 | CS76541 |
| Litva | Litva | CS76543 |
| Lm-2 | Lm-2 | CS76545 |
| Mammo-1 | Mammo-1 | CS76365 |
| Mammo-2 | Mammo-2 | CS76364 |
| Me-0 | Me-0 | CS76549 |
| Mer-6 | Mer-6 | CS76414 |
| Mh-0 | Mh-0 | CS76550 |
| Mitterberg-1-181 | Mitterberg-1-181 | CS76354 |
| Mnz-0 | Mnz-0 | CS76552 |
| Neo-6 | Neo-6 | CS76560 |
| Nie1-2 | Nie1-2 | CS76402 |
| Np-0 | Np-0 | CS76563 |
| Nw-0 | Nw-0 | CS76564 |
| Ob-0 | Ob-0 | CS76566 |
| Or-0 | Or-0 | CS76568 |
| Ove-0 | Ove-0 | CS76569 |
| Pi-0 | Pi-0 | CS76572 |
| Pla-0 | Pla-0 | CS76573 |
| Qar-8a | Qar-8a | CS76581 |
| Qui-0 | Qui-0 | CS76417 |
| Ragl-1 | Ragl-1 | CS76583 |
| Rd-0 | Rd-0 | CS76584 |
| RLD-1 | RLD-1 | CS76588 |
| Rome-1 | Rome-1 | CS76590 |
| Rovero-1 | Rovero-1 | CS76351 |
| Rue3-1-31 | Rue3-1-31 | CS76406 |
| Seattle-0 | Seattle-0 | CS76598 |
| Sei-0 | Sei-0 | CS76559 |
| Shigu-1 | Shigu-1 | CS76375 |
| Shigu-2 | Shigu-2 | CS76374 |
| Sij-1 | Sij-1 | CS76379 |
| Sij-2 | Sij-2 | CS76380 |
| Sij-4 | Sij-4 | CS76381 |
| Slavi-1 | Slavi-1 | CS76419 |
| Star-8 | Star-8 | CS76400 |
| Stepn-1 | Stepn-1 | CS76378 |
| Stepn-2 | Stepn-2 | CS76377 |
| Timpo-1 | Timpo-1 | CS76424 |
| TueSB30-3 | TueSB30-3 | CS76403 |
| TueV-13 | TueV-13 | CS76407 |
| TueWa1-2 | TueWa1-2 | CS76405 |
| Vezzano-2.2 | Vezzano-2.2 | CS76350 |
| Voeran-1 | Voeran-1 | CS76352 |
| Yeg-1 | Yeg-1 | CS76594 |
