## Supplemental Table 2 for "HTL/KAI2 signalling substitutes for light to control plant germination"

**Table S2. Supplemental Table of Gene Expression Data**

**This dataset contains the common genes in Ara-1:KAI2OX and KAI2OX+KAR2:KAI2OX comparisions, whose q-value < 0.05**

| LIGHT | KAI2OX | Ara-1 | *Striga* | | |
|---|---|---|---|---|---|
| **TAIR** | **log2 expression (24 hr)** | | **log2 (16h)** | **Gene name** | **Annotation** |
| AT1G76570 | -2.4 | -2.0 | -0.4 | *LHCB7* | Chlorophyll A-B binding protein |
| AT4G14690 | 7.8 | 3.6 | 3.5 | *ELIP2* | Chlorophyll A-B binding protein |
| AT4G17600 | 3.6 | 2.7 | 0.9 | *LIL3.1* | Chlorophyll A-B binding protein |
| AT1G44575 | 9.0 | 6.2 | 9.0 | *PSBS* | Chlorophyll A-B binding protein |
| AT1G29930 | 1.6 | 2.1 | -1.0 | *LHCB1.3* | Chlorophyll A-B binding protein |
| AT1G29910 | 3.6 | 3.2 | 1.3 | *LHCB1.1* | Chlorophyll A-B binding protein |
| AT4G10340 | 2.4 | 3.6 | 11.4 | *LHCB5* | Chlorophyll A-B binding protein |
| AT3G08940 | -3.9 | -2.1 | 1.1 | *LHCB4.2* | Chlorophyll A-B binding protein |
| AT3G47470 | 1.6 | 1.3 | 1.5 | *LHCA4* | Chlorophyll A-B binding protein |
| AT4G08920 | -0.4 | -0.4 | -0.5 | *CRY1* | cytochrome |
| AT5G24850 | 0.8 | 1.7 | 0.1 | *CRY3* | cytochrome |
| AT1G26260 | 2.9 | 2.1 | -0.8 | *BHLH76* | cytochrome interating protein |
| AT5G38430 | 12.7 | 10.0 | 3.5 | *RBCS-1B* | Rubisco |
| AT5G38420 | 6.7 | 4.6 | 4.4 | *RBCS-2B* | Rubisco |
| AT5G38410 | 7.7 | 5.4 | 1.9 | *RBCS3B* | Rubisco |
| AT1G67090 | 3.0 | 2.7 | NA | *RBCS-1A* | Rubisco |
| AT1G32060 | 5.3 | 4.3 | 5.3 | *PRK* | Phosphoribulokinase |
| AT3G61470 | 4.7 | 3.4 | 5.3 | *LHCA2* | photosystem I light harvesting complex gene 2 |
| AT1G31330 | 4.8 | 3.3 | 2.7 | *PSAF* | photosystem I subunit F |
| AT1G52230 | 7.3 | 4.2 | 1.8 | *PSAH2* | photosystem I subunit H2 |
| AT1G08380 | 6.0 | 6.1 | 7.3 | *PSAO* | photosystem I subunit O |
| AT1G19150 | 6.2 | 4.0 | 1.5 | *LHCA6* | photosystem I light harvesting complex gene 6 |
| AT1G67740 | 6.5 | 4.8 | 3.4 | *PSBY* | photosystem II BY |
| AT3G27690 | 6.3 | 7.3 | 2.4 | *LHCB2.4* | photosystem II light harvesting complex gene 2.3 |
| AT4G15510 | 1.9 | 1.3 | 0.6 | *PPD1* | Photosystem II reaction center PsbP protein |
| AT2G30570 | 1.4 | 1.4 | 2.8 | *PSBW* | photosystem II reaction center W |
| AT5G23120 | 1.7 | 0.6 | 0.4 | *HCF136* | photosystem II stability/assembly factor |
| AT3G50820 | 3.5 | 2.0 | 3.1 | *PSBO2* | photosystem II subunit O-2 |
| AT4G21280 | 5.7 | 4.4 | 3.3 | *PSBQ1* | photosystem II subunit QA |
| AT3G21055 | 4.7 | 4.1 | NA | *PSBTN* | photosystem II subunit T |
| AT5G11450 | 0.9 | 0.4 | 0.9 | *AT3G55610* | photosystem II reaction center PsbP protein |
| AT5G27390 | 1.7 | 0.8 | 1.2 | *AT5G27390* | photosystem II reaction center PsbP family protein |
| AT3G56650 | 2.2 | 1.7 | 0.9 | *PPD6* | photosystem II reaction center PsbP family protein |
| AT3G50330 | 7.0 | 4.9 | 0.8 | *HEC2* | bHLH DNA-binding protein |
| AT1G02340 | 5.2 | 7.8 | NA | *HFR1* | bHLH DNA-binding protein |
| AT2G20180 | -1.8 | -1.1 | -1.8 | PIF1 | bHLH DNA-binding protein |

| GA | KAI2OX | Ara-1 | *Striga* | | |
|---|---|---|---|---|---|
| **TAIR** | **log2 expression (24 hr)** | | **log2 (16h)** | **Gene name** | **Annotation** |
| AT1G15550 | 7.5 | 5.1 | -2.7 | GA3OX1 | GA3 oxidase 1 |
| AT1G80340 | 11.3 | 8.4 | NA | GA3OX2 | GA3 oxidase 2 |
| AT4G25420 | 0.7 | 2.1 | 11.6 | GA20OX1 | GA20 oxidase 1 |
| AT1G30040 | -1.7 | 0.6 | 0.3 | GA2OX2 | GA2 oxidase 2 |
| AT4G09600 | -8.3 | -2.9 | -3.5 | GASA3 | GAST1 protein homolog |
| AT5G15230 | 6.3 | 4.1 | 14.0 | GASA4 | GAST1 protein homolog |
| AT1G74670 | 4.0 | 5.7 | 3.0 | GASA6 | GAST1 protein homolog |
| AT5G14920 | 2.9 | 2.4 | 4.1 | GASA14 | xyloglucan hydrolase |
| AT2G06850 | 6.3 | 4.2 | 8.3 | XTH4 | xyloglucan hydrolase |
| AT5G13870 | 5.9 | 3.7 | 8.4 | XTH5 | xyloglucan hydrolase |
| AT4G03210 | 5.9 | 4.1 | 3.2 | XTH9 | xyloglucan hydrolase |
| AT3G44990 | 4.9 | 4.9 | 9.9 | XTH31 | xyloglucan hydrolase |
| AT1G10550 | 6.5 | 5.8 | 2.3 | XTH33 | xyloglucan hydrolase |
| AT2G20750 | 2.1 | 3.7 | -0.7 | EXPB1 | expansin |
| AT5G05290 | 4.0 | 4.5 | 10.4 | EXPA2 | expansin |
| AT2G37640 | 5.5 | 4.8 | 6.4 | EXPA3 | expansin |
| AT2G40610 | 2.7 | 4.8 | 6.0 | EXPA8 | expansin |
| AT5G02260 | 4.4 | 5.1 | 1.8 | EXPA9 | expansin |
| AT3G03220 | 1.3 | 0.5 | 0.7 | EXPA13 | expansin |
| AT2G03090 | 3.9 | 4.8 | 1.3 | EXPA15 | expansin |
| AT4G38210 | 2.9 | 2.4 | 3.5 | EXPA20 | expansin |
| AT4G12390 | 2.3 | 1.6 | -1.8 | PME1 | pectin methylesterase |
| AT3G14310 | 5.4 | 5.1 | 6.7 | PME3 | pectin methylesterase |
| AT4G25260 | 3.9 | 4.4 | 0.6 | PMEI7 | pectin methylesterase |
| AT4G02330 | 3.3 | 3.6 | 9.4 | PME41 | pectin methylesterase |
| AT4G33220 | 5.1 | 4.0 | 1.7 | PME44 | pectin methylesterase |

| TAIR | log2 expression (24 hr) | | log2 (16h) | Gene name | Annotation |
|---|---|---|---|---|---|
| AT5G45340 | 4.3 | 2.8 | 2.0 | **CYP707A3** | Cytochrome P450 |
| AT3G24220 | -2.8 | -2.1 | 0.4 | **NCED6** | ABA biosynthesis dioxygenase |
| AT1G78390 | -4.8 | -1.3 | 5.3 | **NCED9** | ABA biosynthesis dioxygenase |
| AT5G67030 | -4.3 | -2.0 | 0.6 | **ABA1** | ZEP |
| AT1G77450 | -2.5 | -2.1 | 1.3 | **NAC032** | NAC domain protein |
| AT1G52890 | -3.0 | -1.1 | 1.2 | **NAC019** | NAC domain protein |
| AT1G79270 | -2.3 | -1.8 | -2.6 | **ECT8** | evolutionarily conserved C-terminal domain |
| AT2G22430 | 0.7 | 0.9 | -0.4 | **ATHB-6** | homeobox protein |
| AT3G61890 | 6.6 | 5.8 | 0.7 | **ATHB-12** | homeobox protein |
| AT5G05410 | -2.4 | -1.0 | -0.7 | **DREB2A** | DRE-binding protein |
| AT3G11020 | -4.8 | -2.3 | -1.9 | **DREB2B** | DRE-binding protein |
| AT2G40340 | -3.4 | -1.5 | -1.6 | **DREB2C** | integrase-type DNA-binding protein |
| AT1G75490 | -7.8 | -2.4 | -1.9 | **DREB2D** | integrase-type DNA-binding protein |
| AT5G18450 | -7.5 | -2.6 | -1.9 | **DREB2G** | integrase-type DNA-binding protein |
| AT5G11590 | 4.2 | 4.9 | 0.2 | **DREB3** | integrase-type DNA-binding protein |
| AT1G29390 | 3.3 | 1.8 | 1.3 | **COR413IM2** | cold regulated protein |
| AT2G15970 | -1.4 | -1.2 | 1.2 | **COR413PM1** | cold regulated protein |
| AT3G50830 | 0.6 | -0.4 | 1.3 | **COR413PM2** | cold regulated protein |
| AT5G42900 | -5.5 | -2.5 | NA | **COR27** | cold regulated protein |
| AT1G20440 | 5.5 | 4.6 | NA | **COR47** | cold regulated protein |
| AT3G51810 | -7.1 | -2.6 | -1.4 | **EM1** | stress induced protein |
| AT2G40170 | -8.0 | -3.5 | -1.2 | **EM6** | stress induced protein |
| AT1G70800 | -6.2 | -3.1 | 0.4 | **CAR6** | Ca-dependent lipid-binding protein |
| AT1G70810 | -5.8 | -2.3 | 6.9 | **CAR7** | Ca-dependent lipid-binding protein |
| AT1G70790 | -2.3 | -2.1 | 3.5 | **CAR9** | Ca-dependent lipid-binding protein |
| AT2G36270 | -2.6 | -1.2 | -2.0 | **ABI5** | ABA insensitive 5 |
| AT1G49720 | -3.1 | -2.0 | 1.0 | **ABF1** | ABA response element binding factor |
| AT4G34000 | 5.7 | 3.3 | -1.7 | **ABF3** | ABA response element binding factor |
| AT3G19290 | -3.4 | -1.8 | -0.4 | **ABF4** | ABA response element binding factor |

| ETHYLENE | KAI2OX | Ara-1 | *Striga* | | |
|---|---|---|---|---|---|
| | log2 expression (24 hr) | | log2 (16h) | Gene Name | Annotation |
| AT5G51690 | -0.3 | -0.4 | -1.1 | **ACS12** | AC synthase |
| AT1G01480 | -9.0 | -3.5 | -0.4 | **ACS2** | AC synthase |
| AT4G26200 | 5.1 | 4.1 | 5.7 | **ACS7** | AC synthase |
| AT2G19590 | 3.9 | 4.9 | -0.9 | **ACO1** | AC oxidase |
| AT1G62380 | 1.7 | 2.1 | 0.8 | **ACO2** | AC oxidase |
| AT2G05710 | 2.2 | 2.1 | 1.1 | **ACO3** | AC oxidase |
| AT1G05010 | -2.4 | -1.6 | 2.5 | **ACO4** | AC oxidase |
| AT1G66340 | -0.5 | -0.4 | 1.7 | **ETR1** | Ethylene receptor |
| AT3G23150 | 0.7 | 1.3 | 3.6 | **ETR2** | Ethylene receptor |
| AT2G40940 | 2.9 | 3.2 | 1.9 | **ERS1** | Ethylene receptor |
| AT1G04310 | -4.8 | -1.3 | 1.2 | **ERS2** | Ethylene receptor |
| AT3G04580 | 1.4 | 1.3 | 3.6 | **EIN4** | Ethylene receptor |
| AT5G44790 | -3.2 | -1.6 | 0.0 | **RAN1** | Cu transporter |
| AT3G20770 | -2.4 | -0.9 | -1.3 | **EIN3** | Ethylene transcription factor |
| AT5G10120 | -1.0 | 0.4 | 0.9 | **EIL4** | Ethylene transcription factor |
| AT1G73730 | -1.5 | -0.8 | -0.7 | **EIL3** | Ethylene transcription factor |
| AT1G12980 | 4.5 | 3.9 | 7.5 | **ESR1** | Ethylene response factor |
| AT1G24590 | 4.8 | 5.7 | 9.4 | **ESR2** | Ethylene response factor |
| AT1G53170 | 2.8 | 1.8 | -0.4 | **ERF8** | Ethylene response factor |
| AT1G21910 | 3.4 | 2.6 | 3.4 | **ERF012** | Ethylene response factor |
| AT1G44830 | 6.2 | 4.4 | 0.4 | **ERF014** | Ethylene response factor |
| AT2G44940 | 4.6 | 2.9 | -2.6 | **ERF034** | Ethylene response factor |
| AT3G60490 | 6.4 | 4.3 | 4.4 | **ERF035** | Ethylene response factor |
| AT1G77200 | 3.6 | 2.8 | 0.2 | **ERF037** | Ethylene response factor |
| AT4G27440 | 5.8 | 4.4 | 2.1 | **PORB** | Protochlorophyllide oxidoreductase B |
| AT5G54190 | 3.3 | 2.8 | 5.2 | **PORA** | Protochlorophyllide oxidoreductase A |
